# High-throughput sequencing of unmethylated DNA (UMR-seq) reveals extensive epigenetic variation in the hypomethylome of diverse plant species

**DOI:** 10.64898/2026.10.29.762724

**Authors:** Vanessa Putland, Hannah Robinson, Judith I. M. Eglitis-Sexton, Guilherme V Yoshikawa, Jessica Hintzsche, Haylie Andrews, Stacey Andersen, Sohye Yoon, Jingyang Tong, Leroy M. Mangila, Akshya Sree, Silvia Zanini, Maximilian Schmidt, Yan Zhang, Rod J. Snowdon, Agnieszka Golicz, Kai P. Voss-Fels, Lee T. Hickey, Peter A. Crisp

## Abstract

The unmethylated regions (UMRs) in a genome can reveal the location of genes and gene regulatory loci. However, discovering UMR variation across and between entire genomes and disentangling the contribution of genetics and epigenetics to traits are challenging because they require separate and costly whole-DNA methylome and whole-genome genetic analyses. In many species this is prohibitively expensive for even a small number of samples, curtailing analysis on a population or breeding scale. Here we introduce UMR-seq, a method for the specific capture of UMRs from a genome and simultaneous sequencing of genetic variants in a single economical and scalable assay. UMR-seq charts the hypomethylome, revealing that it is highly enriched with functionally important regulatory loci. While UMRs are highly stable between vegetative tissues in plants, we find they are dynamic in the endosperm and vary substantially between accessions. Adaptable to diverse species, using this method we uncover a positive correlation between gene number in a species and UMR content, reflecting gene regulatory space in a genome. At 1/50^th^ of the conventional cost of methylome sequencing in wheat, UMR-seq enables simultaneous population scale analysis of genetic and epigenetic variation in the functional genomic space. We reveal that in a panel of 187 elite wheat cultivars, profiling UMR variation improved genomic prediction across population groups, and UMR-based association mapping identified trait associations that were largely distinct from those detected with SNPs, providing new opportunities to understand trait biology and to enhance predictive breeding.

## INTRODUCTION

Decoding the information stored in nucleic acids has been transformative to our understanding of life and inheritance. Beyond the DNA bases A, T, G and C, heritable information can be stored in modified bases, such as 5-methylcytosine, commonly known as DNA methylation ^1,2^. Information can also be encoded in chromatin modifications and chromatin structure. Epigenetics concerns heritable phenotypic variation occurring independent of genetic sequence differences ^3^. In plants, flowering time, flower morphology, ripening, fruitset, biomass, yield, architecture, pigmentation and leaf morphology are examples of valuable agronomic traits where beneficial phenotypes have been caused by heritable epigenetic variation (‘epialleles’) without changes to DNA sequence ^4^. In both plants and animals, there is evidence of extensive epigenetic variation between members of the same species, ^5^ or even the same variety, ^6^ and these epigenetic differences can be associated with trait variation. DNA methylation is near ubiquitous in plant genomes, with many contiguous regions displaying consistently high levels of DNA methylation across different plant cells, tissues and organs. A much smaller fraction of most plant genomes displays a consistent lack of DNA methylation. Genomic regions that lack DNA methylation, named <u>U</u>n<u>m</u>ethylated <u>R</u>egions (UMRs), provide extremely useful information for decoding a plant genome because they can predict genes and, importantly, loci enriched for cis-regulatory elements ^7^. In turn, regulatory regions containing cis-regulatory elements facilitate the intricate regulation of gene expression underpinning the complexity of life, as well as being enriched in genetic variants underlying agronomic traits ^8,9^. For these reasons, regulatory regions have also emerged as key targets for genome engineering approaches for plant improvement ^10–12^. High-throughput DNA sequencing technologies have significantly accelerated our understanding of the genome and epigenome over the past two decades. In plants, sequencing 5-methylcytosine has been a principal method to extract nucleotide-level epigenetic information from DNA. Likewise, resolving the chromatin architecture of a locus has been a fruitful approach to identify potential cis-regulatory regions through sequencing combinations of chromatin accessibility, chromatin interactions, histone modifications and DNA methylation ^13,14^. Nevertheless, using DNA methylation profiling to identify UMRs has emerged as arguably the most productive and straightforward method for rapid, genome-wide discovery of candidate regulatory regions in a genome ^7,15^ (**Figure 1a**).

**Figure 1:**
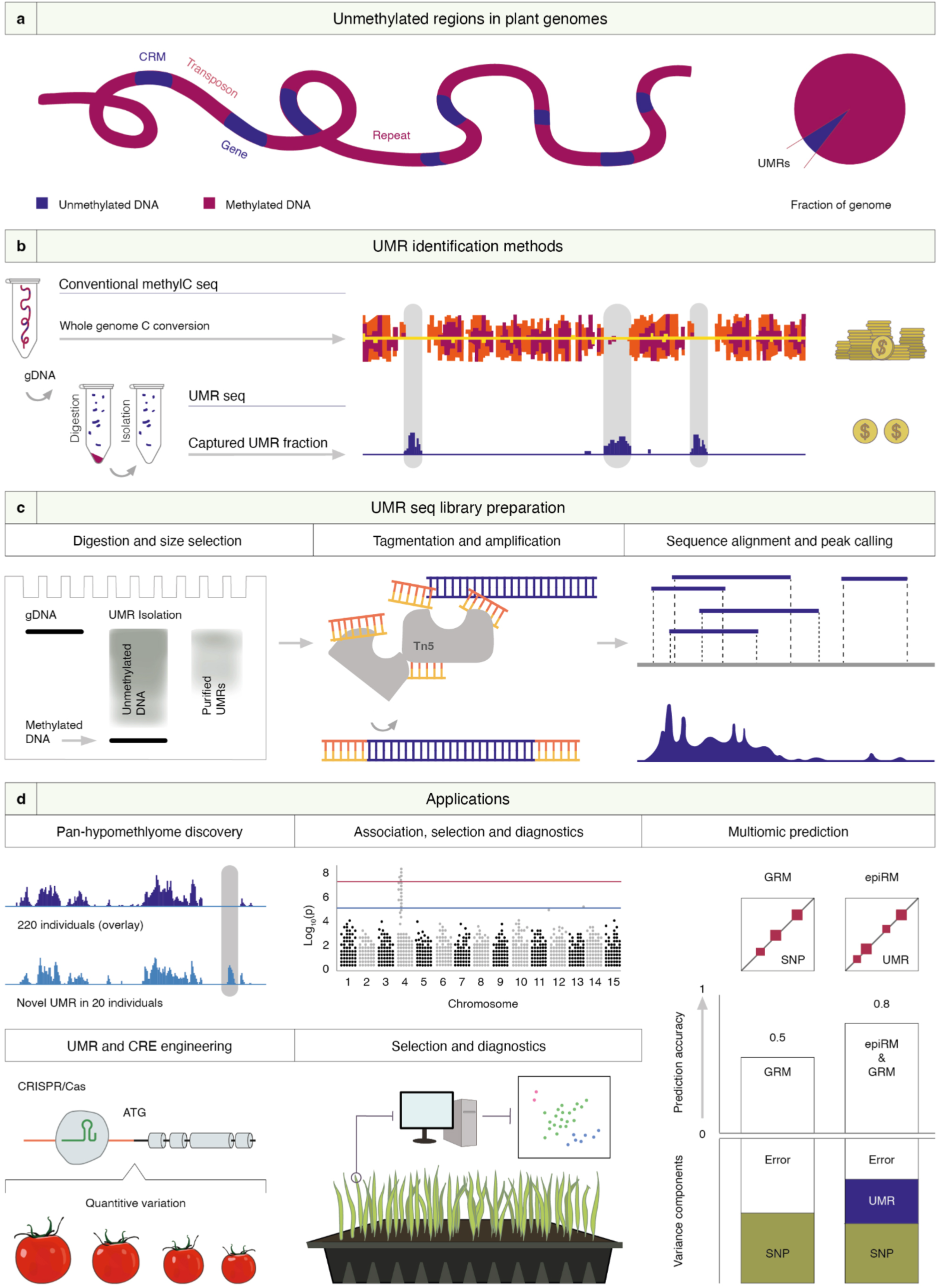
UMR-seq technology overview and applications. **a**, Unmethylated regions (UMRs) mark genes and gene regulatory regions known as cis-regulatory modules (CRMs) that contain cis-regulatory elements (CREs) in plant genomes. In most plant genomes, UMRs comprise a small fraction of the total genome. **b**, In conventional workflows, UMRs are identified using a whole genome methylC-seq profiling strategy that typically uses a C-to-T base conversion approach, followed by whole genome sequencing, which is very costly for plants with large genomes; in contrast, UMR-seq precisely isolates and separates the UMRs from a genomic DNA (gDNA) sample, and only the UMRs are sequenced, generating the same UMR information as conventional methods at a fraction of the cost owing to the dramatically reduced sequencing requirements. **c**, UMR-seq libraries are prepared by isolating UMRs via enzymatic digestion of the genomic DNA with cytosine methyl-dependent restriction endonucleases MspJI and FspEI, followed by SPRI-bead size selection to separate the small methylated fragments generated by digestion from the larger unmethylated regions excised from the methylated gDNA. An example electrophoresis gel visualisation of the isolation and purification steps is depicted. Following UMR isolation, sequencing libraries are prepared using Tn5 enzymatic tagging with adapter sequences (Illumina Nextera workflow), followed by sequencing and genome alignment of the reads, which correspond to the unmethylated regions in a reference genome. UMR-seq produces coverage data (similar to ChIP-seq or ATAC-seq data types) amenable to peak-calling and differential abundance analysis. **d**, The reduced cost of UMR-seq compared to conventional approaches opens the door to larger scale studies such as pan-hypomethylome generation for improved genome annotations for discovery and engineering of gene regulatory elements; UMR-marker based associations (e.g. epi-Genome Wide Association Studies), selection and diagnostics; and integration of UMR information into multi-omic prediction methods to improve prediction accuracy for plant improvement (GRM genetic relationship matrix; epiRM epigenetic relationship matrix).

For crop species with moderate- to large-sized genomes (e.g. >500 Mbp), commonly used high-throughput sequencing approaches are not capable of cost-effective, simultaneous, genome-wide discovery of genetic, epigenetic and cis-regulatory information. These inefficiencies also make profiling populations intractable, stifling the discovery and incorporation of epigenetic information into crop improvement programs for future food security. Conventional next-generation sequencing technologies accurately identify the canonical bases A, T, G and C. To distinguish 5^m^C (and other modifications) from unmodified bases, multiple base-conversion plus sequencing strategies (methylC-seq) have been developed, including enzymatic conversion as employed in enzymatic-methyl sequencing (EM-seq) ^16^, and the long-standing gold-standard approach, whole genome bisulfite sequencing (WGBS) ^17^. However, the conversion of C-to-T (via U) creates an increased computational burden, an ambiguity in the identification of genetic C-to-T polymorphisms (a common mutation), and the low sequence diversity after conversion reduces mappability of reads, specifically affecting regions with few methylated cytosines, such as UMRs ^18^. More significantly, a substantial shortfall of existing approaches is the necessity to profile an entire genome in order to identify UMRs. Yet, UMRs are only a small percentage of most genomes, for example, 3-5% in barley or 6% in wheat (**Figure 1a-b**); meaning that the >94% of the data corresponding to methylated regions is discarded in such workflows. Nevertheless, the most significant limitation of current approaches is the economics of profiling large numbers of individuals. For example, WGBS for hexaploid bread wheat costs many thousands of dollars (>USD$3,500) for just a single sample at adequate coverage due to the 17 Gbp size of the genome. Third generation long-read sequencing methodologies, such as Nanopore and Pacbio, now offer possibilities for simultaneous decoding of modified and unmodified bases ^19^, but also come at a high price point. A suite of more economical alternatives have been developed, ranging from targeted sequence capture to untargeted restriction enzyme-based genome reduction strategies that use methylation sensitive enzymes, including epiRADseq, bsRADseq and epiGBS ^20–25^. Sequence capture methods are useful once loci of interest are identified, but are unsuitable for the discovery of new epialleles. Conventional genome reduction strategies also suffer notable limitations ^24^. Their untargeted nature means much of the data falls within uninformative regions of the genome, which in total is only a very small fraction of a genome (e.g. < 0.0001% ^24^). Using these strategies, only a small number of cytosines can be profiled per locus; however, changes in multiple cytosines across a region (termed a differentially methylated region) are of higher biological significance; plus these approaches can still require base-conversion. Therefore, alternative economical methods are needed for effective discovery and comprehensive characterisation of UMR variation.

Some decades ago, an approach named methyl-filtration was developed as a gene enrichment strategy to enable more efficient shotgun sequencing of the genic portion of plant genomes ^26–28^. A bacterial host strain possessing a gene encoding the methylation specific restriction enzyme McrBC was used to eliminate, and thereby ‘filter’, clones containing methylated repetitive DNA. This method enriches for clones with gene sequences, which typically have low or no methylation. Methyl-filtration was applied to analyse and sequence cloned contigs depleted of densely methylated DNA ^26–33^. Here, inspired by this prior concept, we develop and optimise a new method named <u>U</u>n<u>m</u>ethylated <u>R</u>egion-<u>seq</u>uencing (UMR-seq). This method is used to reveal the total unmethylated space in a genome, including both genic and intergenic regions, and enables unparalleled discovery of epigenetic variation across entire populations. Utilising a new combination of enzymes, UMR-seq is capable of high resolution targeted capture of unmethylated DNA to provide a readout of UMRs, plus simultaneous detection of genetic variants such as single nucleotide polymorphisms (SNPs). Critically, this approach is versatile, scalable and highly economical owing to the targeted isolation of UMRs. The cost is proportional to the UMR content of a genome, which is correlated with the total number of genes. For example, UMR-seq library preparation and sequencing currently costs about a tenth less than methylC-seq in maize, or around 1/30^th^ for barley, both diploid species of average gene content but different genome sizes; for the largest genome we processed, hexaploid wheat, methylC-seq costs approximately 50 times more than UMR-seq (**Figure 1b-c**). We demonstrate the application of UMR-seq in nine different plant species, then profile a breeding panel of elite wheat cultivars to characterise population-scale UMR variation and to evaluate its use in genomic prediction and association mapping. This population-scale analysis, costing a fraction of conventional methylC-seq (a 98% cost saving), shows that UMR data can improve genomic prediction across population groups and identify trait associations that could unlock greater biological insight.

## RESULTS

### The development of UMR-seq

Considering the functional significance of the unmethylated portion of plant genomes, which we refer to as the ‘hypomethylome’, we designed a method for highly efficient capture of unmethylated DNA. Methylation-sensitive restriction endonucleases have been employed in DNA methylation analyses for many years ^34–37^. For most restriction enzymes, the presence of DNA methylation inhibits their activity and they cannot bind or cleave methylated sites; however, a rarer class of enzymes are methylation-dependent, these are specifically active only on methylated recognition sequences. For example, the endonuclease McrBC specifically recognises methylated DNA at each of two half-sites of the form R^m^C (R = G or A), which can be separated by ∼40-3000 bp and cleaves about 30 bp from the R^m^C site ^38,39^ (**Supplementary Fig. 1a**). Inspired by methyl-filtration, which uses *E. coli* clones harbouring the McrBC restriction system ^40^ to deplete methylated DNA in shotgun libraries ^26–33^, we were motivated to develop a method to precisely isolate UMRs (contiguous regions of unmethylated DNA) from purified genomic DNA samples. We wanted to capture UMRs regardless of their genomic context, including small UMRs (several hundred base pairs) surrounded by dense methylation. In this respect, McrBC has several shortcomings, including producing fragments of hundreds-to-thousands of base pairs; such fragments may not accurately define borders between methylated and unmethylated DNA. McrBC also has a limited (rare) recognition sequence (G/A^m^C), which is less effective against the non-CG methylation prevalent in plant genomes, and McrBC requires an extended incubation time. Therefore, we sought alternative enzymes that could more precisely isolate unmethylated DNA from purified genomic DNA. MspJI and FspEI belong to a family of modification-dependent restriction endonucleases that cleave methylated DNA at a fixed distance from the modification site ^41,42^. MspJI recognises 5^m^C in the context ^m^CNNR and cleaves 16 bp 3’ to the methylated sites. This means that for symmetric 5^m^C sites (e.g. CG and CHG; H = A, T or C), the two methylated half sites are recognised by a pair of MspJI enzymes, leading to excision of a 32 bp fragment of methylated genomic DNA (**Supplementary Fig. 1b**). FspEI works in the same manner as MspJI and has the recognition sequence C^m^C (**Supplementary Fig. 1c**). Previously, methylated DNA excised by these enzymes has been used to profile the methylated portion of a genome ^41^. However, here we optimised a method to isolate and sequence the undigested and unmethylated DNA fraction, which is the basis for UMR-seq (**Figure 1b-c**). In this method, genomic DNA is treated with MspJI and FspEI (Methods) to cleave methylated DNA into ∼32 bp fragments, leaving a higher molecular weight unmethylated DNA fraction containing fragments of contiguous unmethylated DNA (**Figure 1c**). The higher molecular weight unmethylated fraction is then separated from the lower molecular weight methylated fraction using a paramagnetic bead isolation to exclude any fragments <100 bp (**Figure 1c**, **Supplementary Fig. 2**). The isolated, unmethylated DNA can then be prepared for next generation sequencing. In this workflow, we use the Illumina Tn5-based Nextera XT DNA Library Preparation Kit; however, this approach would be equally compatible with other NGS library preparation methods suitable for analysis of DNA fragments or amplicons (**Figure 1c**). Libraries can then be sequenced using standard NGS sequencing methodologies to generate coverage maps of the unmethylated portion of a genome, also known as the hypomethylome (**Figure 1b-c**). Because this method precisely captures the small unmethylated fraction of a genome, it is considerably cheaper than conventional UMR profiling methods, opening the door to applications that require larger scale profiling of plant populations, such as discovery and engineering of conserved CREs from a pan-methylome, epi-marker based selection or diagnostics, and epi-genomic prediction for plant improvement (**Figure 1d**).

Considering that MspJI and FspEI do not recognise all possible ^m^C sites in a genome (and some regions lack C altogether), some regions of methylated DNA will be recalcitrant to cleavage into fragments <100 bp during the UMR-seq purification step. Such regions need to be excluded from further analyses because the methylation status cannot be evaluated using UMR-seq, and could result in false positive UMR identification. Therefore, we performed *in silico* digestion modelling (Methods) to determine the fraction of a genome that can be effectively profiled using UMR-seq. We compared using MspJI or FspEI individually, or both combined (**Supplementary Table 1**). Using the 2.4 Gbp maize genome as an example, we found 2,731 and 227,808 regions were recalcitrant to cleavage into <100 bp fragments by MspJI and FspEI, respectively. When combined, only 1124 regions could not be eliminated by MspJI + FspEI. These 1124 regions represent only 0.0079% (168,162 bp) of the maize genome, and are recorded as a ‘blacklist’ of regions to be excluded from analysis of UMR-seq data. In fact, 99.61% of the black-listed regions represent simple sequence repeats (e.g. [TATAG]_n_) that do not contain the recognition motif of the enzymes; only 6 of the black-listed regions were not composed of repetitive sequences. The blacklist regions are recalcitrant to methylC-seq analysis as well (**Supplementary Fig. 3**) and >90% are not detectable above background in UMR-seq data (**Supplementary Fig. 4**), but it is still best practice to exclude them. Similar results were obtained for sorghum, barley and wheat, with 0.02%, 0.05% and 0.003% of each genome, respectively, black-listed. For each species analysed by UMR-seq we have generated a blacklist of regions to be excluded from analysis.

### UMR-seq precisely maps the hypomethylome

We generated different levels of digestion using the UMR-seq enzymes to determine the optimal range of digestion conditions required to remove methylated DNA for maize, barley and wheat (**Supplementary Figs. 5-8**). For all three species, when using 1x enzyme concentration (5 units total), digesting 0.5 or 1.0 μg of DNA for 120 minutes provided the highest enrichment for known UMRs and isolated regions with the lowest DNA methylation, based on EM-seq data. These conditions also yielded the highest Fraction of Reads in Peaks (FRiP), ranging from 70-85%. ATAC-seq data has similar qualities to UMR-seq data and the ENCODE recommended FRiP for ATAC-seq is >30%. Therefore, depending on sample availability, 0.5 to 1.0 μg of DNA digested for 120 minutes was selected as optimal for further analyses of maize, barley and wheat. We also compared UMR-seq (MspJI + FspEI) to digestion with McrBC followed by sequencing (McrBC-seq) (**Supplementary Fig. 9**) and found that McrBC-seq did show promising enrichment for unmethylated regions; however, UMR-seq had superior performance. Replicate UMR-seq samples were highly correlated (Pearson’s r > 0.96), including replicates of the same genotype and when comparing to a different genotype (**Figure 2a**). To determine the optimal read depth, we deeply re-sequenced the maize leaf UMR-seq libraries, obtaining up to 200 million reads from which discrete sets were randomly subsampled (**Supplementary Fig. 10**). We found that sequencing UMRs to 10 million reads struck an optimal peak/cost ratio. Peak counts plateaued between 10 and 100 million reads, although the UMR space continued to increase in this interval, possibly by increasing peak width.

**Figure 2:**
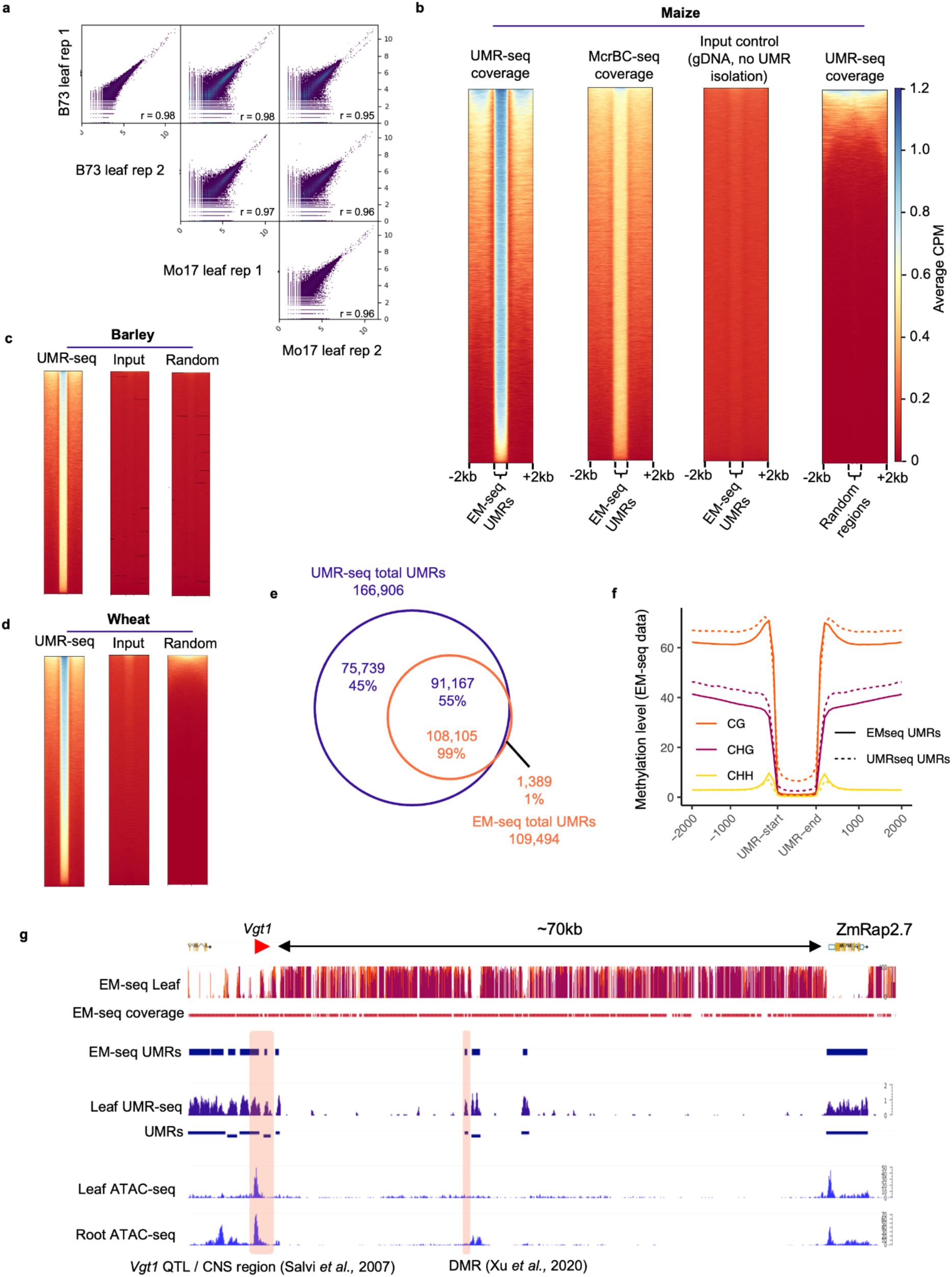
UMR-seq implementation and benchmarking. **a**, Pearson correlation for read counts over UMR-seq UMRs between maize replicates and genotypes (B73 and Mo17). **b,** Read coverage for maize B73, visualised over known EM-seq UMRs, comparing from left to right: UMR-seq, McrBC-seq, gDNA whole genome sequencing (negative ‘input control’ made by omitting the UMR isolation enzymes), and UMR-seq read coverage over randomly selected genomic regions. **c**, Read coverage for barley Morex, and (**d)** wheat Chinese spring, over known methylC-seq UMRs and controls. **e**, UMR-seq UMRs were identified using epic2 and their genomic location was compared to methylC-seq UMRs in maize B73; co-located regions indicated by the overlapped region in the Venn diagram have >20% reciprocal overlap in genomic coordinates. **f**, The methylation profile of UMR-seq UMRs determined by EM-seq. **g**, Example UMR-seq profile at the *Vgt1* locus (B73 AGPv4 8:135969172-135969381), UMRs (pink highlighted regions) that mark the *Vgt1* enhancer element ^47,49^ and the domestication enhancer ^48^ are efficiently captured using UMR-seq.

UMR-seq data generated for maize, barley, and wheat, was evaluated against methylC-seq data generated from the same DNA samples for maize and barley (including the highest quality reference UMR profile generated for barley to date), and against a published methylC-seq dataset for wheat ^43^, representing the first UMR analysis in wheat. For each species, we first identified UMRs using conventional methylC-seq data then compared these ‘known’ UMRs to UMR-seq data. We found that UMR-seq data coverage was highly specific and highly enriched over the methylC-seq-identified UMRs in all species (**Figure 2b-d**). For example, in maize, less than 1% (1,389) of the EM-seq UMRs were not detected using UMR-seq data (**Figure 2e**). Next, using a peak-calling approach, we identified UMRs in each species using UMR-seq data, and determined the methylation levels for these regions using methylC-seq data. We found that these UMR-seq UMRs were highly depleted of methylation in each context (CG, CHG and CHH) in maize (**Figure 2f**; **Supplementary Fig. 5**), barley (**Supplementary Fig. 7**) and wheat (**Supplementary Fig. 8**). UMR-seq identifies significantly more UMRs relative to methylC-seq (**Figure 2e**) and our analysis of the UMRs that were unique to UMR-seq in maize suggests this is caused by multiple factors, but mainly due to finer resolution of UMR-seq, partial or sporadic methylation coupled with the inherently differing bioinformatic approaches. UMRs unique to UMR-seq were very sparse in cytosine data from EM-seq (**Supplementary Fig. 11a**). Sparsity was not due to a complete lack of cytosines in the UMRs, although these regions were relatively depleted of CG and CHG, and more frequently contained only CHH sites (**Supplementary Fig. 11b**). Unique UMR-seq UMRs also tended to be enriched in intergenic gene distal regions (>2 kbp from a gene) (**Supplementary Fig. 11c**). Low sequence diversity after conversion theoretically reduces mappability of EM-seq reads in unmethylated regions, which could be exacerbated in intergenic, repeat-rich regions of the maize genome. However, we observed very little difference in EM-seq read coverage at shared UMRs compared to UMR-seq unique UMRs (**Supplementary Fig. 11d-e**). The standard EM-seq UMR analysis pipeline omits UMRs if the region only contains CHH sites, so including these CHH-only regions in the analysis captured a further 20,010 of the UMR-seq UMRs originally considered unique (**Supplementary Fig. 11f**). UMR-seq unique UMRs also had on average a lower peak confidence (epic2 q-score), which would be expected for regions only unmethylated in a subset of cells producing a lower UMR signal (**Supplementary Fig. 11g**). Another factor contributing to differences between EM-seq and UMR-seq UMRs is that our methylC-seq pipeline only includes UMRs 300 bp or larger^44^ but UMR-seq can detect smaller UMRs (**Supplementary Fig. 11h-i**). For example, where two 200 bp UMRs are separated by sporadic methylation and not classed as a UMR in EM-seq, UMR-seq would count one or two peaks depending on the rolling window peak-calling settings used (**Supplementary Fig. 11j**). Some of the residual methylation detected at UMR-seq UMRs, in particular in the CG context (**Figure 2f**), would be expected for certain classes of partially methylated regions including those caused by heterozygous UMRs. Likewise, if a subset of the leaf cells profiled were methylated while the majority of cells were unmethylated at a locus, this would be expected to give rise to a signal in UMR-seq data despite the mixed cell population. Using DNA from hybrid maize samples, we tested this hypothesis and found that UMR-seq can detect heterozygous UMRs, not detected using the conventional UMR analysis method for methylC-seq data (**Supplementary Fig. 12c**). If desired, the majority of UMR-seq unique UMRs could be removed from a UMR-seq analysis by filtering using a minimum CG/CHG density threshold combined with a size filter, pre- or post-peak calling, to retain a set of UMRs with greater similarity to standard EM-seq UMRs (**Supplementary Fig. 12k**). We also found that peak callers had differing propensity to identify partially methylated regions; MACS2 ^45^ favoured regions that lacked any methylation signal in EM-seq data, while epic2 ^46^ detected more partially methylated regions (**Supplementary Fig. 12a-b, d**), suggesting that the choice of peak caller may depend on the biological question being addressed. Together, these factors mean that UMR-seq detected a larger number of UMRs in total, compared to methylC-seq (**Figure 2e**).

UMR-seq identified known non-coding regions and cis-regulatory elements. For example, the *Vgt1* enhancer element upstream of *Rap2.7* ^47^, which affects flowering time in maize, and the characterised UMR enhancer that arose after maize domestication from teosinte ^48^, are both marked by UMRs and efficiently identified using UMR-seq (**Figure 2g**). The hypomethylome for maize and other species can be browsed online at https://crisplab.genome.edu.au/apollo.

### The hypomethylome is enriched with functionally important regulatory loci

Unmethylated regions are known to be highly enriched in functionally important regions of the genome ^7^. Using functional data available for maize, we investigated whether the UMR-seq hypomethylome is also able to efficiently capture these functionally important regions. We found enrichment for genes, accessible chromatin regions (ACRs, from ATAC-seq), conserved non-coding sequences (CNS), and transcription factor binding sites (TFBS, from ChIP-seq) (**Figure 3a**), as well as characterised CREs (**Figure 2g**). Over annotated genes and their proximal regions (± 2 kbp), UMR-seq coverage showed strong enrichment at transcription start and end sites (**Figure 3a**). Of the 39,498 annotated genes, we found that 33,835 genes and their proximal regions overlapped with UMRs. Looking at promoter regions alone (3 kbp upstream gene annotations), 30,876 overlapped with UMRs. The analysis was repeated for publicly available ATAC-seq data ^7,14,50^, resulting in a pattern similar to that of UMR-seq coverage, though with a sharp signal at transcription start sites and a less pronounced peak at transcription end sites. In leaf, most of the ACRs (94%) overlapped with UMRs. Notably, 94-97% of ACRs from root, ear and grain also overlapped with leaf UMRs, with 50-75% overlap between vegetative tissue ACRs (**Figure 3b**), indicating that regions with potential to become accessible chromatin are marked by UMRs in the leaf hypomethylome even when they are inaccessible in this tissue. Furthermore, UMR-seq coverage was depleted over transposable elements, as expected of actively suppressed genomic elements (**Figure 3a**). Conversely, coverage was highly enriched at CNS (**Figure 3a**). CNS are cis-regulatory sequences conserved across phylogenetic taxa, with some ancestral sequences reportedly conserved since before the genesis of the seed plant clade ^51^. Of the 394,900 CNS annotated in the maize genome, 281,000 (71%) CNS - the majority - overlap with 37,810 (22%) of the leaf UMRs (p < 0.05) (**Figure 3c-d**). Similarly, using a published ChIP-seq dataset for 104 TFs expressed in leaf tissue ^52^, we found that 87% of transcription factor binding sites overlap with 53% of the leaf UMRs (p < 0.0005), aligning with our previous findings ^7^, and an overlap greater than for conventional methylC-seq UMRs (**Figure 3e**-**f**). Given the close link between UMRs and ACRs, it is unsurprising that our findings are concordant with the conclusions of a comprehensive ACR study in maize and other species by Lu *et al*. ^53^. These results show that the hypomethylome is heavily enriched for functional loci relevant across different plant cells, tissues and organs, and UMR-seq can distill this information by profiling a single easy-to-sample organ, such as a leaf.

**Figure 3:**
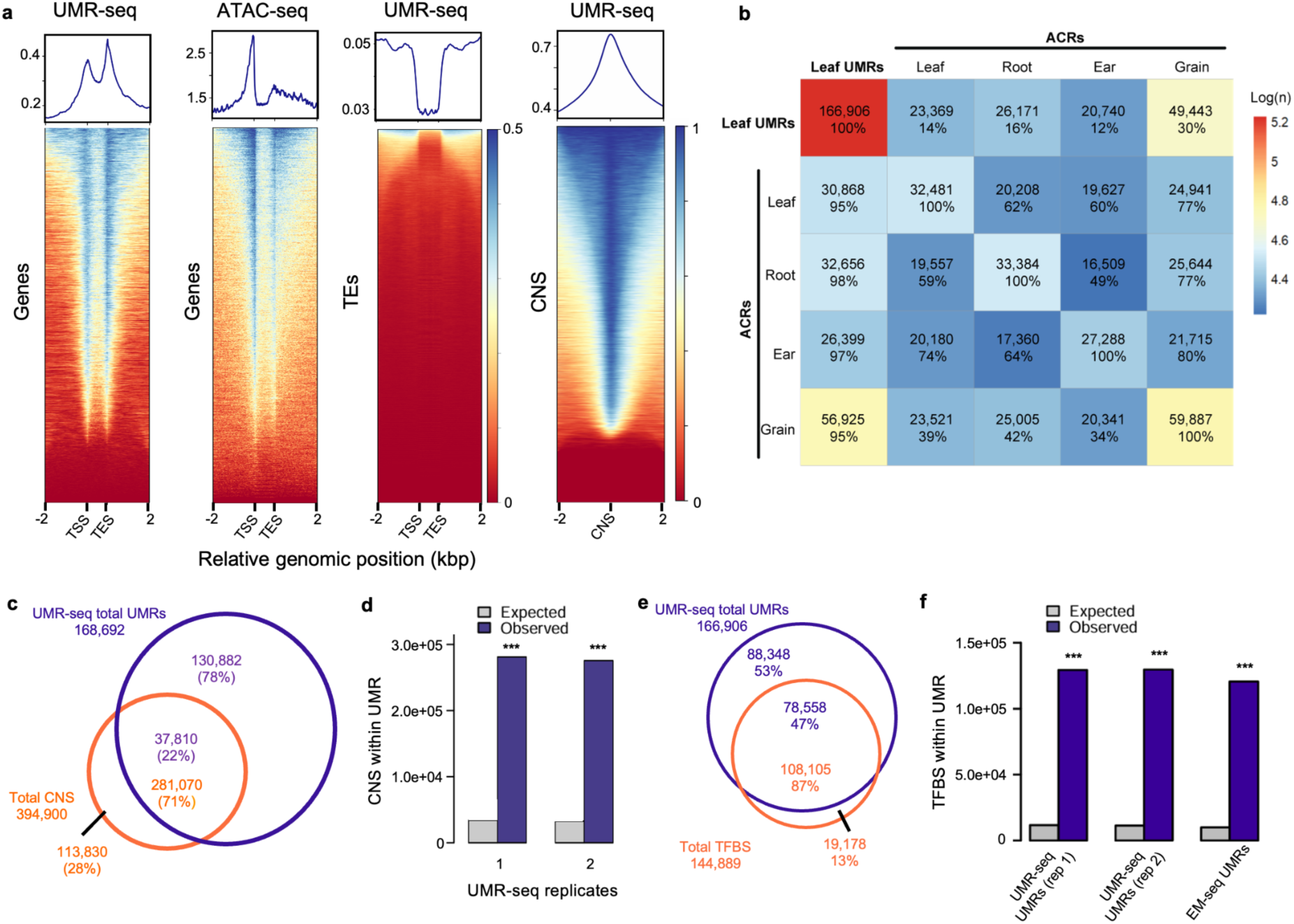
Enrichment of functional elements in UMRs in maize. **a,** Enrichment rates of chromatin features in UMR-seq profiles: genes, transposable elements (TE), genes in accessible chromatin regions (ACR), and conserved non-coding sequences (CNS). ACR regions were profiled using ATAC-seq ^7,14,50^. The representative enrichment profiles are centred on the feature of interest, displaying 2kb upstream of the transcription start site (TSS) and 2 kb downstream of the transcription end site (TES); or 2 kb flanking either side of the centre of the CNS. **b,** Proportion of leaf UMRs from UMR-seq overlapping with ACRs from ATAC-seq of samples from leaf, root, ear and grain, and overlap of ACRs between the various tissues. **c**, Proportion of CNS overlapping with UMRs. **d**, Observed enrichment of CNS ^54^ within UMRs from two UMR-seq replicates compared to the overlap expected by random chance (P = 0.0002) based on 5,000 permutations of randomised regions. The analyses shown in (**c**) and (**d**) were carried out using the B73 v5.0 reference genome. **e**, Proportion of UMRs overlapping with transcription factor bindings sites (TFBS). **f**, Expected and observed overlap between transcription factor binding sites (TFBS) and UMRs from two replicates of UMR-seq, compared to UMRs from enzymatic methyl-seq, P = 0.0002 between the three expected and observed comparisons, based on 5,000 permutations of randomised regions. *** is p < 0.001.

### UMR-seq is effective across diverse plant species without modification

In principle, the UMR-seq method is species agnostic, but the relative benefit scales with genome size. Therefore, we applied UMR-seq to nine different plant species to demonstrate the broad applicability, while testing genome sizes ranging from 430 Mbp to 17,000 Mbp (**Supplementary Fig. 13a**) across rice, grapevine, sorghum, tomato, soybean, maize, *Nicotiana benthamiana*, barley and wheat. In contrast to the gold-standard methylC-seq, where UMRs are computationally identified from whole genome datasets by filtering for regions that contain less than 10% methylation at cytosines, UMR-seq enzymatically isolates UMRs, leading to a huge saving in sequencing costs, proportional to genome size. First, *in silico* digestion modelling was conducted to predict recalcitrant regions and the estimated analysable proportion of each genome was >99.9% (**Supplementary Table 2, Supplementary Fig. 13a**). Dicots had a slightly higher number of recalcitrant regions, likely related to lower GC content (**Supplementary Fig. 13b,** 33-37% GC for dicots compared to 42-45% for the monocots). We set out to compare the UMRs resulting from UMR-seq versus those from methylC-seq **(Figure 4a-b)**. For each species, we overlayed normalised UMR-seq read coverage, and control gDNA read coverage, over known UMR regions identified by methylC-seq. For maize, barley and grapevine we generated methylC-seq data from the same DNA used for UMR-seq, while for other species we reanalysed published methylC-seq data (**Supplementary Table 3**). We found strong alignment between regions with high coverage in UMR-seq and the known UMR regions, and observed that UMR-seq was highly enriched at known UMRs in each of the species when compared to the coverage in the matched gDNA undigested control **(Figure 4a)**. UMR-seq identified more UMRs than methylC-seq in most species, with the exception of grapevine, and the majority of UMRs identified by methylC-seq (58-99%) were also detected by UMR-seq **(Figure 4b)**. To investigate the performance of UMR-seq in grapevine further, we sequenced seven paired UMR-seq and EM-seq samples observing 21-43% of EM-seq UMRs uniquely identified in EM-seq (**Supplementary Fig. 14**). We attribute this to the inconsistent per-sample peak identification by the algorithm (despite highly similar coverage profiles). This is most likely due to the large proportion of unmethylated DNA in grapevine, potentially in combination with high heterozygosity affecting background estimation and UMR peak detection. Merging the individual UMR-seq peak profiles from these clones (all were the same genotype) did not increase the total number of UMRs (108,054 compared to the individual range of 66,322–116,621), but gave an improved overlap and peak detection, reducing the EM-seq unique UMRs from 21–43% to 11%. The greatest total difference in UMRs between sequencing approaches were identified in barley and wheat with UMR-seq identifying ∼200,000 more UMRs than methylC-seq in each **(Figure 4b)**. UMR-seq generally identified more total base pairs of unmethylated DNA (UMR space) than methylC-seq, including in barley, wheat, maize, *N. benthamiana*, tomato and sorghum. The greatest difference was seen in wheat, with 241.56 Mbp further UMR space identified by UMR-seq compared to methylC-seq. In soybean, grapevine and rice, methylC-seq found a greater amount of UMR space **(Supplementary Table 4)**. The average UMR length was longer in five species in UMR-seq data, including wheat, *N. benthamiana*, tomato, sorghum, grapevine, compared to methylC-seq, while UMRs were on average shorter in the other species, with the greatest difference seen in soybean where UMR-seq UMRs were over 500 bp shorter on average relative to methylC-seq **(Supplementary Table 4)**. These results demonstrate that UMR-seq reliably detects UMRs across a range of plant species.

**Figure 4:**
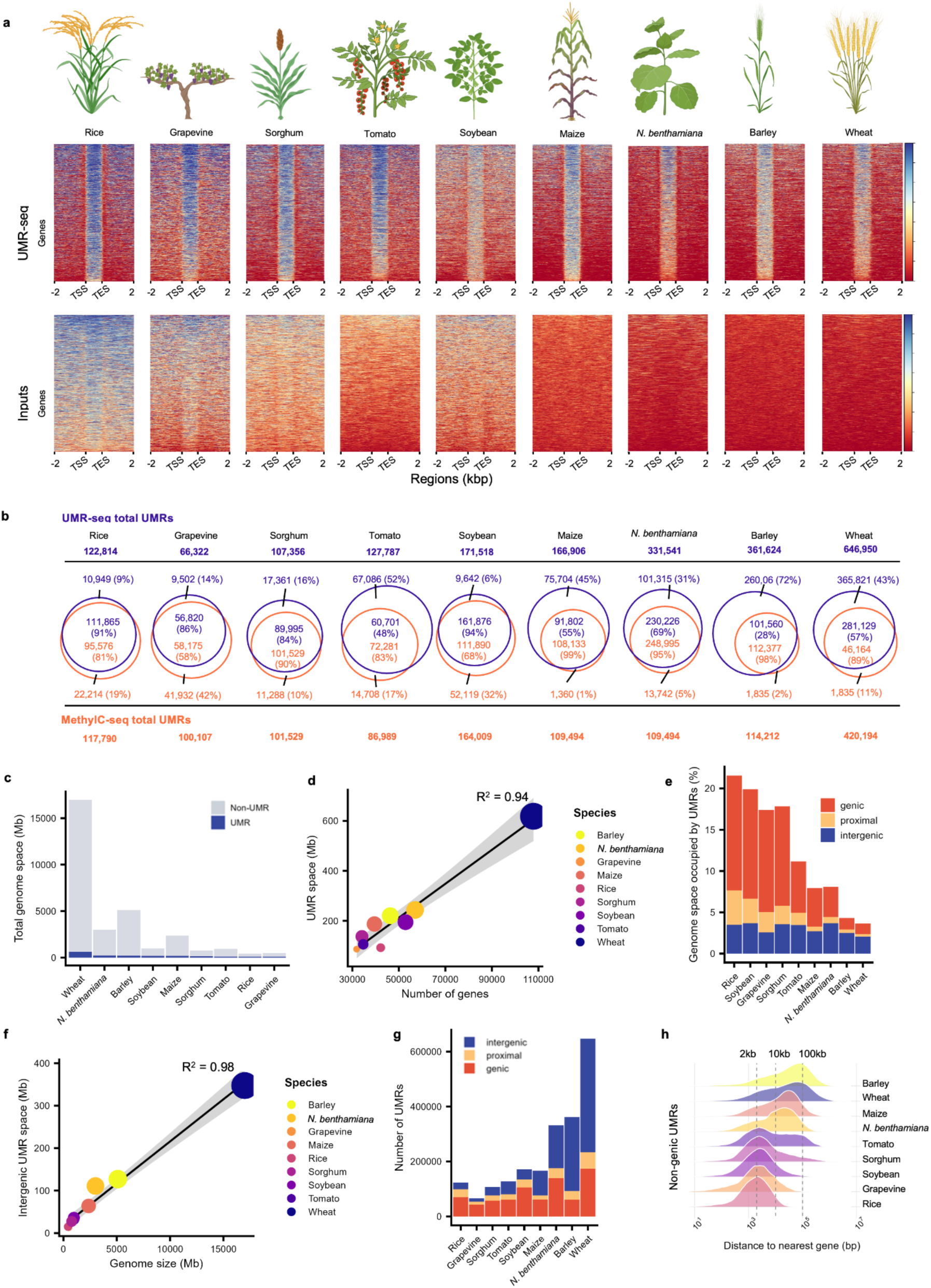
UMR-seq application in multiple plant species. **a**, UMR-seq signal across methylC-seq unmethylated regions. Normalised UMR-seq signal is shown for methylC-seq unmethylated regions (1 kbp) by signal strength across nine plant species in UMR isolated (UMR-seq) and control (gDNA) samples, ordered by genome size (from left to right, pictorial graphics represent rice, grapevine, sorghum, tomato, soybean, maize, *Nicotiana benthamiana* (*N. benthamiana*), barley and wheat. Species UMR-seq and gDNA control samples were bioinformatically processed using identical parameters; colour bars indicate relative signal strength, with colour scale values omitted due to species specific maximum signal values. The x-axis is centred on genic regions. Negative ‘input control’ was made by omitting the UMR isolation enzymes, therefore consisting of undigested genomic DNA. **b**, Number of UMRs identified by UMR-seq (purple) vs methylC-seq (orange), and overlap of UMRs detected between the two approaches. **c**, Proportion of the genome captured by UMR-seq (dark blue) versus total genome size of the corresponding reference genome (light gray), illustrating how UMRs represent a fraction of a genome. **d**, Relationship between UMR space (Mbp) and number of genes reveals a strong correlation between the two variables, confirmed by a coefficient of determination (R^2^) of 0.97. The trend line was calculated by linear regression, with a 95% confidence interval shaded in gray. Trend line equation is included inset. Spot size is determined by the genome size, and colour is determined by the species (distributed in the spectrum by alphabetic order). **e**, Percentage of genomic space occupied by UMRs from UMR-seq for each plant species studied. UMRs were categorised by their position relative to the nearest gene: genic (red, located within gene boundaries), proximal (yellow, within 2 kbp of the nearest gene), and intergenic (blue, more than 2 kbp away from the nearest gene). The distribution of UMRs in each genome shows that the proportion of genic and proximal UMRs tends to shrink as genome size increases, while intergenic UMRs remain roughly as a constant proportion of each genome regardless of species. Species are arranged on the x-axis by ascending genome size. **f**, Investigating the relationship between the genome space occupied by intergenic UMRs (Mbp) versus the genome size (Mbp) revealed an almost perfect proportional correlation, with a coefficient of determination (R^2^) of 0.98. Spot size is determined by genome size, spot colour is determined by species. **g**, Number of UMRs per category in each species. With genomes ordered by ascending size on the x-axis, it is apparent that number of genic UMRs is linked to ploidy (therefore, gene number and not genome size), and the number of intergenic UMRs increases with genome size. **g,** Density plots of non-genic UMRs (proximal and intergenic) relative to their distance to the nearest gene (bp) show that the density peaks shift away from the gene as the genome size increases. Figure components Created in BioRender.

For each species, we compiled the identified UMRs from UMR-seq into a hypomethylome and determined the fraction of each genome composed of UMRs (**Figure 4c**). Previously using methylC data we observed that the total base pairs of genomic space (in Mbp) occupied by UMRs was consistent between diploid grass species despite large variation in genome size ^7^. With UMR-seq we can now investigate this relationship in more detail, and we found that the UMR space is strongly correlated with the number of genes in a genome (R^2^ = 0.97, **Figure 4d; Supplementary Fig. 15).** Accordingly, the UMR space in hexaploid wheat (107,891 genes, 618 Mbp UMR space) is approximately triple to the UMR space of barley (46,294 genes - 221 Mbp UMR space), a closely related diploid species also in the Triteceae tribe. Considering the relationship between gene number and total UMRs (or unmethylated genomic space), we developed a linear model to calculate the target read depth in any species based on our calculation of 10 million reads for maize (**Supplementary Fig. 10**), the model can also be used to predict the UMR space (Mbp) in a genome and cost per hypomethylome (y = 0.0068x-126.8, **Supplemental Table 5**). Genome size is also correlated with the number of genes, consequently genome size is strongly correlated with UMR space (**Supplementary Fig. 15**). Across the nine species, UMRs covered between 3-22% of genome space (**Figure 4e**). In smaller genomes (<1,500 Mbp), the proportion of UMRs was 10-22%, whereas in larger genomes (>2,000 Mbp), UMRs covered closer to 3-8% (**Figure 4e**), indicating that although UMR space is positively correlated to genome size, the proportion of genome occupied by UMRs is inversely proportional to genome size (R^2^ = 0.46, **Supplementary Fig. 15**). We then compared the spatial distribution of UMRs across the various species’ genomes and classified them into three groups based on position relative to genes: genic UMRs (i.e., found within the boundaries of a gene annotation), proximal UMRs (i.e., located within 2,000 kbp of the nearest gene), and intergenic UMRs (i.e., located more than 2,000 kbp away from the nearest gene) (**Figure 4e, Supplementary Table 6**). We found that while the proportion of the genome occupied by proximal and genic UMRs reduced as the genome size increased, from 14% of the rice genome to 1% in wheat (**Figure 4e**), the absolute genic and proximal UMR space tended to increase with genome size (proximal R^2^ = 0.71, genic R^2^ = 0.82, **Supplementary Fig. 15, Supplementary Table 6**). The results from UMR-seq show that the genome coverage by intergenic UMRs remained comparatively stable across species at 2-4% of the genome (**Figure 4e**), meaning that intergenic UMR space increases with genome size (R^2^ = 0.98), as does the number of intergenic UMRs (**Figure 4f-e, Supplementary Table 6**). The number of genic UMRs is not linked to genome size, instead appearing to be associated to ploidy number ^55–59^ and, consequently, to gene number (**Figure 4g, Supplementary Table 6**). This contrasts with the reduction in intergenic UMR fraction as the genome increases reported by Crisp *et al.* ^7^ from analysis of WGBS data, indicating that UMR-seq has improved capability to capture intergenic UMRs over the capability of WGBS. Lastly, we investigated the location of proximal and intergenic UMRs relative to the nearest gene, and found that non-genic UMRs (intergenic + proximal) in smaller genomes were concentrated within 2-10 kbp of the closest gene. As genome sizes increased, the non-genic UMR population density peaks shifted towards 100 kbp (**Figure 4f**). Given that the number of proximal UMRs also increases with genome size (**Supplementary Table 6, Supplementary Fig. 15**), the most likely explanation for the change in location distribution is that the increased complexity of larger genomes leads to the transposition or rewiring of UMRs (possibly via the action of transposable elements), or the emergence of novel UMRs in intergenic regions.

### UMR-seq enables highly cost-effective, replicated differential methylation analysis

UMR-seq data can be summarised across the genome into a read-count or coverage format, closely resembling other data types such as ATAC-seq, RNA-seq or ChIP-seq. Accordingly, this data is compatible with a variety of differential analysis methods developed for epigenomic data types, including differential count or differential peak analysis. We implemented several different approaches to compare UMRs between samples, including a pipeline built around csaw ^60^ (Methods) for the identification of differentially unmethylated regions using biological replicates. We have used this tool to investigate conserved and stable UMRs between cultivars of a species, organs, or tissues, adding to our understanding of epigenetic variation and regulation. We first set out to identify differential UMRs in maize, comparing different tissue types, including endosperm and leaf, and the two genotypes: B73 and Mo17. Peaks were called in the UMR-seq profile by epic2 and the results were compared on leaf tissue between the two genotypes. There was over 75% overlap in peak location, indicating strong conservation across genotypes (**Figure 5a**). The algorithm csaw ^60^ was used to perform a statistical comparison of UMR-seq signal in sliding windows across the genome to identify regions of differential UMR abundance (which we term D-UMRs) between the two sample groups, finding a total of 65,385 D-UMRs (**Figure 5b**). Initially, this was surprising as this number is similar to the total number of unique peaks in each sample (75,361, **Figure 5a**). While there is extensive genomic variation between the two genotypes ^61^, a WGBS analysis showed that UMRs are depleted in polymorphic regions ^62^, so a large number of D-UMRs is unlikely to be due to polymorphism or poor sequence mapping. However, D-UMRs do not necessarily directly correspond to the ‘UMRs’ identified by running a peak caller such as epic2, D-UMRs can represent a change in UMRs abundance among cells (heterogeneous sample), a change in UMR size (the edge of a UMR), and D-UMRs can be smaller than an epic2-UMR because the csaw sliding window is 100 bp (multiple D-UMRs per UMR). Previously, using WGBS data, we identified 107,178 UMRs in B73, of which ∼16,300 (15.3%) were unique to B73 and absent from Mo17 ^62,63^. Thus, the UMR-seq csaw D-UMR analysis pipeline discovers more differential UMRs compared to methylC-seq analyses.

**Figure 5.**
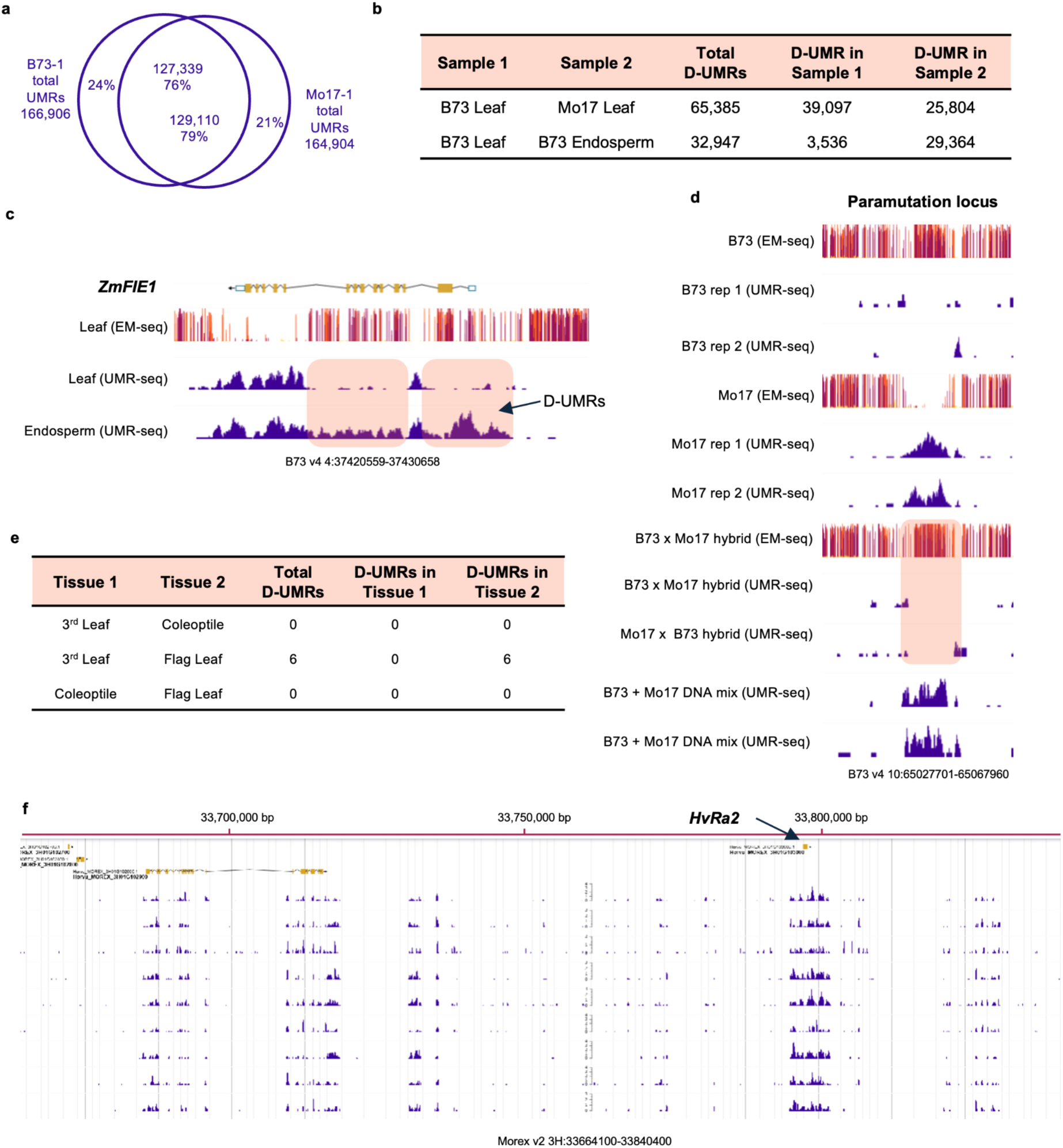
Identification of differential UMRs (D-UMRs) between genotypes and tissue types in maize and barley. **a**, Venn diagram showing the intersection and unique UMR-seq peaks between samples of the maize genotypes B73 and Mo17. **b**, Statistical comparison of D-UMRs between divergent samples: from leaf tissue between two genotypes (B73 and Mo17), and between leaf and endosperm tissue from the genotype B73. “D-UMRs in Sample X” are exclusive to that sample. **c**, Visualisation of D-UMRs found between maize leaf and endosperm samples, on the imprinted gene *FERTILIZATION INDEPENDENT ENDOSPERM1* (*ZmFIE1*, *Zm00001d049608*), the parental allele of which is methylated ^65^. Orange shading marks the D-UMR. **d**, Example of a D-UMR associated with a paramutation, showing EM-seq and UMR-seq profiles of the parents B73 and Mo17, and of the reciprocal offspring exhibiting the paramutation. **e,** Statistical comparison of D-UMRs between three barley tissue types in background RGT Planet, tissues sampled and compared were third leaf, flagleaf and root. **f**, Comparison of barley UMR-seq profiles near the gene *RAMOSA2* (*HvRa2*, *Horvu_MOREX_3H01G103000*) between three tissues, three biological replicates each. Window size is approximately 188 kbp in the Morex v2 genome.

Next, two tissues were chosen for comparison, including the endosperm, which due to its hypomethylated state ^64^ was expected to yield a large number of D-UMRs relative to the leaf. There were a substantial number of D-UMRs (32,947) identified between leaf and endosperm, although only around half the number of D-UMRs called between genotypes (**Figure 5b**). As an example, we looked at the methylation profile of the imprinted gene *FERTILIZATION INDEPENDENT ENDOSPERM1* (*FIE1*, *Zm00001d049608*), the maternal allele of which is demethylated in the endosperm ^65^ (**Figure 5c**). In leaf, *FIE1* is methylated at most exons, as shown by the presence of coverage peaks in EM-seq and absence thereof in UMR-seq. Conversely, in the endosperm sample, UMR-seq reads cover the whole gene, indicating that it is unmethylated (**Figure 5c**) and suggesting that UMR-seq can likely detect allele-specific UMRs because maternal allele demethylation underpins *FIE1* imprinting. In addition to detecting heterozygous methylation states, the UMR-seq analysis pipeline is able to detect D-UMRs in lineages, as exemplified in the paramutation-like pattern described in **Figure 5d**. A paramutation is a phenomenon characterised by the transfer of epigenetic information between alleles (DNA methylation, in this case) that results in a heritable change of expression of one allele ^66,67^. We found a locus unmethylated in genotype Mo17 and methylated in B73, but it was methylated in the offspring of both reciprocal crosses, confirmed by both EM-seq and UMR-seq. This indicates that the positive methylation status of the allele inherited from the B73 parent may have been transferred to the allele inherited from Mo17. Thus, by sequencing the entire hypomethylome, UMR-seq will enable large-scale investigation of paramutations.

To investigate whether the results from maize can be generalised to other grass species, we carried out the same UMR characterisation on three barley tissues/organs (root, third leaf and flag leaf) in the genetic background of cultivar RGT Planet. In contrast to the thousands of D-UMRs detected between maize genotypes, only a few D-UMRs were found between aerial tissues, with six D-UMRs identified between third leaf and flag leaf (**Figure 5e**). This indicates the existence of tissue-specific D-UMRs, albeit a negligible number when compared to the D-UMRs found between maize genotypes (**Figure 5b**). Visualisation of the UMR-seq profiles from the various tissues further supports the conclusion that UMRs are stable across samples originating from a single genotype, enabling reliable comparisons that can be scaled to population-wide analyses (**Figure 5f**). These results show that UMR-seq can be used to detect differentially methylated regions within and between individuals.

### Population-scale UMR-seq reveals a diverse pan-hypomethylome

The utility of UMR-seq at the population level was investigated by applying the technology to a panel of 187 European elite wheat varieties from the BRIWECS panel and previously described by Voss-Fels *et al.*^68,69^. In comparison to the reference hypomethylome of the Chinese Spring IWGSC RefSeq v1.0 genome, the pan-hypomethylome comprises 1.38 Mbp, representing a 141% increase over the reference (**Figure 6a**). This expanded diversity in the marker set provides the basis for population-scale analysis. While UMR profiles were broadly similar between samples (**Supplementary Fig. 16**), many locus-specific differences were observed. To identify polymorphic UMRs on a population scale, we implemented a ‘Consensus UMR’ analysis pipeline used to identify the presence/absence of a UMR (UMR allele, or ‘epiallele’) in each sample (Methods, **Supplementary Fig. 17**).

**Figure 6:**
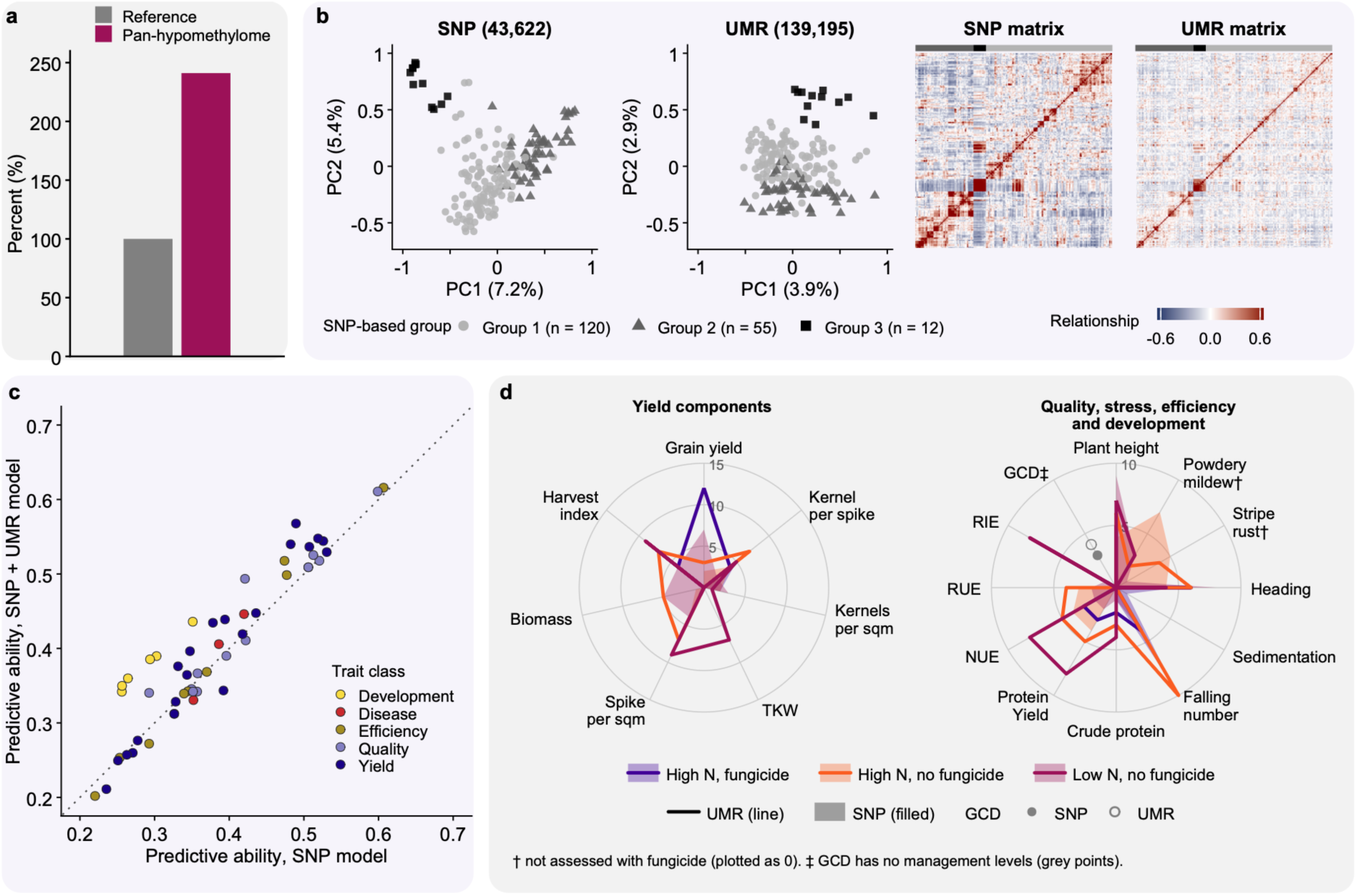
Population-scale UMR-seq in a European elite wheat breeding panel. **a**, the pan-hypomethylome represents a 141% increase in UMR count relative to the reference. **b**, Population structure of the panel from SNP-derived UMR-seq reads (SNP) and UMRs. Left: principal coordinates 1 and 2 of the SNP and UMR relationship matrices. Right: the SNP and UMR (non-orthogonalised) relationship matrices, ordered by dendrogram of the leading ten SNP principal coordinates. Grey shades and symbols in the principal-coordinate plots, and the grey bars above the matrices, show the three dendrogram groups. **c**, Genomic prediction across population groups. X-axis: mean predictive ability of the SNP-only model against the y-axis: SNP-plus-UMR model for 53 trait–management combinations. Points above the dotted 1:1 line (33 of 53) improved prediction with the UMR kernel; colours show trait class (Supplementary Table 7). **d**, Number of significant GWAS associations (FDR q ≤ 0.03) per trait and management for UMRs (lines) and SNPs (filled polygons), for yield components (left) and quality, stress, efficiency and development traits (right); colours show nitrogen and fungicide management. Powdery mildew and stripe rust were not assessed with fungicide (plotted as 0); GCD has no management levels.

### UMR-seq simultaneously provides UMR and SNP markers for population analysis

UMR-seq data from the 187 lines in the wheat panel were used to test whether the technology can be applied at population scale and whether the resulting data can inform predictive breeding and selection decisions, through analyses of population structure, genomic prediction and genome-wide association. Prediction and association analyses covered 53 trait–management combinations spanning yield components, development, grain quality, disease resistance and nitrogen- and radiation-use efficiency. Traits were evaluated in multi-location field trials over two seasons under three nitrogen and fungicide management regimes and analysed as adjusted means across locations and seasons within each management regime^69^. Filtering for minor allele frequency (MAF) > 0.05 and no more than 10% missing calls yielded 139,195 polymorphic UMRs and 43,622 SNPs from the same sequencing reads (hereafter SNPs). SNP-derived estimates of pairwise relatedness were consistent with those from an independent 15K SNP array (off-diagonal r = 0.87; **Supplementary Fig. 18**). UMR markers were distributed across all 21 chromosomes, whereas both SNP datasets showed sparse coverage in chromosome centers and the D subgenome. The distribution of SNPs broadly matched that of SNP-array markers (Spearman ρ = 0.73 for marker counts across 5 Mbp windows; **Supplementary Fig. 19**).

### UMR and SNP variation capture related but partly distinct structure in the wheat panel

SNP and UMR relationship matrices captured shared patterns of relatedness across the panel, with UMR relationships showing weaker separation between population groups (**Figure 6b**). The first two principal coordinates explained 7.2% and 5.4% of SNP variation, compared with 3.9% and 2.9% of UMR variation, respectively, consistent with the limited genetic stratification previously reported for this panel^69^. Ward clustering of the leading ten SNP principal coordinates identified three groups of 120, 55 and 12 lines. These groups overlapped more in UMR coordinate space, although the smallest group remained distinct along the second coordinate. The off-diagonal elements of the two relationship matrices were correlated (**Figure 6b**; r = 0.73), indicating shared structure.

### UMR variation improves prediction across wheat panel population groups

To assess the contribution of UMR variation to genomic prediction, a SNP-only GBLUP model was compared with a two-kernel model incorporating a UMR relationship matrix orthogonalised with respect to the leading 16 SNP principal components in order to largely remove the shared structure. Prediction was evaluated using a leave-one-population-out cross-validation scheme across five population groups defined from SNP relationships, with each group predicted using the other four as the training set. Predictive ability improved in 33 of 53 trait–management combinations (62%; **Figure 6c**; Wilcoxon P = 0.003; **Supplementary Table 7**). Improvements occurred in every trait class and management regime and were most consistent for developmental traits, where predictive ability increased in all six combinations. Gains also increased with trait heritability (Spearman ρ = 0.65, P < 0.001): predictive ability improved in 5 of 18, 13 of 17 and 15 of 18 combinations in the lower (SNP-only h² 0.31–0.71), middle (0.71–0.82) and upper (0.83–0.99) thirds of heritability, respectively. Adding the UMR kernel also reduced prediction bias in 68% of combinations and root mean squared error in 59%, indicating predictions that were better calibrated to, and closer to, the observed values (**Supplementary Fig. 20**).

### UMR-GWAS and SNP-GWAS identify largely distinct trait associations

Genome-wide association studies (GWAS) were performed separately for 132,433 UMRs and 42,905 SNPs with chromosome positions where both analyses used the same lines, traits and association model. At a false discovery rate threshold of q ≤ 0.03, 196 UMR associations were detected across 38 trait-management combinations and 136 SNP associations across 39 combinations (**Figure 6d**; **Supplementary Table 8**). UMR associations outnumbered SNP associations in each management regime, although approximately three times as many UMRs as SNPs were tested. Thirteen UMR associations were in linkage disequilibrium (LD; r² > 0.2) with a significant SNP association for the same trait, compared with 1.3 expected by chance (permutation P = 0.001). The remaining 183 UMR associations were not in LD with a significant SNP association for the corresponding trait. Associations were detected in every trait class (**Figure 6d**). Developmental traits, which showed the most consistent prediction gains, also had the most associations per trait–management combination for both marker types and 3 of the 13 UMR associations in LD with a SNP association were for plant height. UMR associations outnumbered SNP associations for yield, quality and efficiency traits, whereas SNP associations were more frequent for disease traits (**Supplementary Table 7**).

## DISCUSSION

There are limited examples of well-studied DNA methylation epialleles, and most are in model plants with small, simple genomes ^1,2,70,71^. Given the enormous size and complexity of many crop plant genomes, these few examples likely represent only the tip of the iceberg. This knowledge gap principally stems from the prohibitive cost of whole-genome DNA methylation profiling, curtailing widespread discovery and limiting our understanding of the contribution of DNA methylation to trait variation. Here, we have presented the development, optimisation and application of UMR-seq, a cost-effective technological approach to precisely capture and sequence the hypomethylome, and demonstrate that UMR-seq is highly efficient, cheap, reproducible, and scalable. The wheat genome was the largest genome we studied and for wheat methylC-seq-based UMR analysis costs 50 times more than UMR-seq. Thus, UMR-seq enabled a 98% saving on costs. Sequencing costs differ greatly in different parts of the world, so there is significant potential for this savings to be even higher. As an illustrative example, in our hands, library preparation presently costs AUD$33 per sample, giving a total cost per hypomethylome of AUD$53 for maize, for a depth of 10M reads at a rate of AUD$20 per 10M reads, and AUD$93 for wheat for 30M reads, in contrast to the ∼AUD$5,000 required for a methylC-seq-based UMR analysis in wheat.

UMR-seq is a method for the precise capture of the unmethylated regions in a genome. Benchmarking in maize and other species demonstrated that UMR-seq can capture >99.99% of UMRs identified by the conventional approach. While parallels could be drawn to reduced-representation methods, or other low-cost methods employing methylation-sensitive enzymes, UMR-seq differs fundamentally in its principle, because it *enriches* and captures all the UMRs in a genome. For example, the method methylation-sensitive amplified fragment length polymorphism (MS-AFLP/MSAP) exploits the different methylation sensitivity of isoschizomer enzymes that have the same recognition sequence. At CCGG sites HpaII is sensitive to methylation of either C while MspI is only sensitive to methylation at the outer C ^72,73^. The differential display of a band on a gel or sequence read can be used to deduce the methylation status of a specific cytosine site. These methylation-sensitive enzymes have also been used to cleave unmethylated CGs, then the fragments (or fragment ends) analysed by next-generation sequencing to identify unmethylated CGs at the recognition site. This has been termed MRE-seq and can use HpaII (CCGG), Hin6I (GCGC) and AciI (CCGC) ^74^. However, these methods can only reveal the methylation status of a specific cytosine, rather than regional levels, and are sparse due to the genome reduction, subsampling a small fraction of the total cytosines in a genome. Restriction-site Associated DNA sequencing (RAD-seq) ^22^ works by enriching a pseudo-random but reproducible subset of regions of the genome using restriction enzymes, followed by sequencing. epiRADseq (analogous to MRE-seq) is a double-digest RAD-seq (ddRAD) variant in which one of the two enzymes is methylation-sensitive ^21^. bsRAD-seq is also derived from RAD-seq, and involves bisulfite sequence conversion on the subset of regions recovered after genome reduction ^23,24^. Similarly, epiGBS is derived from genotyping-by-sequencing (GBS) and uses a digest for reduced representation followed by bisulfite conversion to measure DNA methylation within fragments ^20,75^. epiGBS digestion with PstI (which is sensitive to CHG methylation) in *Arabidopsis* isolated 2,260 loci, representing ∼0.37% of the genome ^20^. Thus, these restriction enzyme-based approaches are limited by sparse coverage and the enzyme-only methods are restricted to analysis only at a specific recognition site ^35^. While useful for subsampling methylation patterns in a genome, existing methods are underpowered for locus discovery because causal loci can be missed. Likewise genome-wide analysis of UMRs needs contiguous, regional coverage, which these sparse and site-specific methods cannot provide. A clear limitation of UMR-seq is that it cannot (by design) analyse the methylated portion of plant genomes and so cannot decipher contributions of specific methylation contexts or pathways, such as regulation of CG, CHG and CHH. Nevertheless, UMR-seq enables genome-wide comparison of UMRs between samples and within populations, making it an ideal approach for differentially methylated loci discovery and trait association in large genomes.

We have developed several analysis approaches for UMR-seq borrowing from established methods for analogous data types (RNA-seq, ATAC-seq). As is the case for RNA-seq, ATAC-seq and other methods, the choice of analysis pipeline can affect the output, which we observed, for example, when using different peak-callers. Thus, we anticipate ongoing development of UMR-seq analysis pipelines as data for more applications and species is produced. Likewise, we are confident UMR-seq can detect heterozygous UMRs, and further analysis can explore the quantitative resolution.

UMR-seq simultaneously generated a dense set of UMR markers distributed across the wheat genome for the wheat panel, together with SNP markers that recovered patterns of relatedness consistent with an independent SNP array. Both marker types supported standard quantitative-genetic analyses, including multi-kernel genomic prediction and GWAS. Previous studies have explored multi-omics prediction in barley ^76,77^, perennial ryegrass ^78^ and maize ^63^ by combining separate genotyping and omics platforms. By capturing SNP and UMR variation in a single assay, UMR-seq offers a practical route to applying these approaches at the breeding-population scale ^79,80^. Adding UMR variation improved prediction across population groups on average, with gains in every trait class and management regime. Predicting groups excluded from training approximates the practical challenge encountered in breeding when new germplasm enters a breeding program and is less closely related to the training population. The introduction of new genetic diversity in breeding programs is essential for sustaining genetic gain, particularly as genomic selection has the potential to shorten breeding cycles and deplete existing diversity more rapidly. The results suggest that UMR data can complement SNP-based prediction across population structure, motivating further evaluation in newly introduced germplasm and independent breeding populations.

Explicit modelling of genotype-by-environment interactions was beyond the scope of this analysis, which used adjusted means across locations and seasons within each management regime ^69^. DNA methylation can respond to environmental conditions ^81^, raising the possibility that UMR variation captures environment-dependent differences in trait expression. However, analysis of a cross-environment means cannot establish whether UMR-associated effects are consistent across environments or vary with growing conditions. Jointly modelling SNP and UMR effects and their interactions with the environment is therefore an important next step.

UMR-based association analysis identified trait-associated regions that were poorly covered by the tested SNPs. This limits the extent to which the UMR signals can be distinguished from underlying sequence variation. Of the 163 distinct UMRs with significant trait associations, 61% were located in regions with fewer than five tested SNPs within 1 Mbp, highlighting the limited SNP coverage around many of these associations. These regions occurred predominantly in chromosome centers and the D sub-genome, where UMR coverage was extensive but SNPs were sparse (**Supplementary Fig. 19**). UMR associations may therefore reflect methylation variation, sequence variation not represented by the tested SNPs, including structural or presence–absence variants, or both. Although the associations do not establish a functional role for methylation, they provide targets for exploring the biology underlying trait variation. Such insight could extend our understanding of trait genetic architecture and, in turn, strengthen predictive models and selection decisions.

Applying UMR-seq to 187 elite wheat lines shows that the technology can be used at population scale and that the resulting UMR data add value to predictive breeding, improving prediction across population groups. The UMR associations identified here open new opportunities to understand the biology underlying trait genetic architecture. Another exciting future direction is to apply UMR-seq to more diverse panels, where UMR variation could be even higher and there is more power to disentangle the genetic and epigenetic contributions to phenotypes ^5^. Together, these findings suggest that UMR-seq can serve as a beneficial additional data layer for predictive breeding, for informing breeding decisions and a platform for unparalleled investigation of UMR variation and the contribution of epigenetics to traits.

## METHODS

### Plant materials for UMR-seq development and benchmarking across species

Seedlings of barley, maize, *Nicotiana benthamiana*, rice, sorghum, soybean, tomato and wheat **(Supplementary Table 3)** were grown in Conviron growth chambers with UQ23 + Osmocote (16 hours at 24℃ daylight, 60% RH). Samples of young leaf tissue up until six weeks-old-plants were harvested and flash-frozen in liquid nitrogen and stored at -20°C. Leaf samples from grapevine were grown and harvested at Hochschule Geisenheim University, Germany. The European wheat breeding panel and the associated phenotypic data are described in full in Voss-Fels *et al*. ^68,69^. Further details are included in **Supplementary Methods.**

### UMR-seq library preparation

UMR-sequencing samples are listed in **Supplementary Table 9**. Samples for the digestion optimisation were from a single genotype, while each species for the cross species analysis included a minimum of two biological replicates and an undigested, genomic DNA (gDNA) control (**Supplementary Table 9**). Purified and intact DNA was extracted from barley, maize, *N. benthamiana*, rice, sorghum, soybean, tomato and wheat using the Bioline ISOLATE II Plant DNA Kit or Promega Maxwell RSC instrument and the Maxwell RSC PureFood GMO and Authentication kit, and grapevine by Innut PLANT DNA Extraction kit (Analytik Jena). UMR-seq libraries were prepared starting with purified genomic DNA as input **(Supplementary Table 10)** then enzymatically digested with the methylation-dependent restriction enzymes MspJI and FspEI to preferentially deplete methylated DNA and enrich for unmethylated genomic regions. Digestion was performed in a final reaction volume of 30 µl containing 2.5 U each of MspJI and FspEI, with 1x CutSmart buffer and 1x Enzyme Activator. Reactions were incubated first at 37°C for between 90 to 180 minutes followed by enzyme inactivation for 20 minutes at 65°C. The quantity of DNA input and digestion times are dependent on species, where factors such as genome size, methylation levels of the genome and sample purity should be considered (**Supplementary Table 10)**. Larger UMR fragments were then size-selected to separate them from the small, methylated digest fragments (31-32 bp) using a SPRIselect bead ratio of 0.7x to exclude any fragments of 100-200 bp and smaller. The quality of the digestion was analysed by a gel electrophoresis and DNA quantified on a Qubit fluorometer with the 1x High Sensitivity assay (Life Technologies, Eugene, OR, USA), to ensure a complete digestion of methylated DNA occurred and sufficient unmethylated DNA was recovered **(Supplementary Figure 2)**. Purified unmethylated DNA was diluted to 0.2 ng/µl, and 1 ng was used for library preparation. Sequencing libraries were prepared using the Nextera XT DNA Library Prep Kit (**Supplementary Methods**). Libraries were sequenced to 10-20 million reads for the diploid species, 20 million reads for tetraploid *N. benthamiana* and 30 million reads for hexaploid wheat (**Supplementary Table 9**). Subsequently, using ultra-deep sequencing and sub-sampling, we determined 10M reads is optimal for maize and target read depth can be estimated for other species using the total number of genes in a genome (**Supplementary Table 5**).

### UMR-seq read processing and alignment

Reads were processed using the UMR-seq bioinformatic pipeline (available at github.com/pedrocrisp/umrseq). For each species respectively, reads were quality checked by fastqc ^82^ and trimmed by TrimGalore ^83^, then aligned to reference genome **(Supplementary Table 2)** using bowtie2 ^84^. Recalcitrant UMR regions predicted by *in silico digest* (**Supplementary Methods**) in each species were removed prior to read normalisation. Mapped reads were normalised by deepTools bamCoverage (--binSize 10 –normalizeUsing CPM –extendRead) excluding blacklisted regions from *in silico* predictions to enable visualisation in IGV and Jbrowse (**Supplementary Methods**) ^85^. deepTools v3.5.1 source code was modified (following the developers instructions) to disable merging of neighbouring zero bins to enable UMR quantification and comparison in each discrete genomic bin. Normalised UMR-seq read coverage over regions of interest was obtained by deepTools computeMatrix. UMRs were identified and called as peaks from mapped reads by epic2 v3.8 (--bin-size 50) using single-end parameters ^46^ and MACS2 ^45^ using pair-end parameters with median coverage across peaks calculated by Mosdepth ^86^. Peaks were then quality checked by fraction of reads in peaks (FRiP) using deepTools plotEnrichment ^85^ **(Supplementary Table 9)**. The number of UMRs, total UMR space and mean UMR space were calculated from peak files **(Supplementary Table 4 & 6)**.

### MethylC-seq

MethylC-seq samples are listed in **Supplementary Table 3**. UMRs were identified across nine plant species; rice ^87^, grapevine, sorghum, tomato ^88^, soybean ^89^, *N. benthamiana* ^90^, barley and wheat ^59^ utilising in-house and published methylC-seq datasets as previously described ^7^. EM-seq libraries were prepared using the NEB Next Enzymatic Methyl-seq Kit (**Supplementary Methods**). Reads were processed using the methylC-seq UMR bioinformatic pipeline (available at github.com/pedrocrisp/crisplab_epigenomics/tree/master/methylome-s) and described in Eglitis-Sexton *et al*., 2024 ^44^, using the bwameth ^91^ and MethylDackel pipeline variant.

### Functional analysis of UMR-seq

deepTools v3.5.1 ^85^ was used to produce heatmaps for UMR-coverage over methylC-seq data, transposable elements, conserved non-coding sequences (CNS) ^54^, and normalised ATAC-seq read coverage over gene annotations. BEDtools intersect ^92^ was used to quantify the number of both UMR peaks overlapping promoters, genes, and CNS regions and, conversely, CNS regions overlapping the UMR peaks. regioneR v1.44.0^93^ was used to assess the enrichment of CNS regions within UMRs by comparing the observed number of overlaps with the expected distribution generated from 5,000 permutations ^93^. This was repeated for transcription factor binding sites. Details are included in **Supplementary Methods**.

### UMR-genome relationships

Gene content of genomes were determined using GFF3 annotations downloaded from Phytozome (https://phytozome-next.jgi.doe.gov/); except for tomato, barley and wheat that were downloaded from https://solgenomics.net/, https://graingenes.org/GG3/ (via IPK) and https://www.wheatgenome.org/, respectively **(Supplementary Table 3)**. The total number of genes and genome size were compared against total UMR space by linear model in R using the mean UMR space of two biological replicates of each species **(Supplementary Tables 2, 4)**.

### Population UMR analysis in wheat and the pan-hypomethylome

The population scale analysis in wheat used 187 of the 191 elite European winter wheat cultivars from the BRIWECS panel, phenotyped at six locations over two seasons under three management intensities (high nitrogen with fungicide, high nitrogen without fungicide and low nitrogen without fungicide)^69^. Published BLUEs for 53 trait–management combinations covering yield components, development, grain quality, disease resistance and resource-use efficiency were used as phenotypes^69^. The ‘pan-hypomethylome’ in wheat was generated by identifying UMRs per sample using peaks identified by epic2, then merging the peak files together into consensus UMRs. This consensus-UMR pipeline, used for population analysis, was conceived and implemented by AG (available at github.com/pedrocrisp/umrseq) and is described in detail in **Supplementary Fig. 17**. Briefly, per-sample UMRs were merged across samples with bedtools into a consensus set, using the median boundaries of overlapping UMRs. Reads supporting each consensus UMR were counted per sample with featureCounts and normalised to CPM with edgeR ^94,95^. Each UMR was then called present or absent per sample by fitting a kernel density estimate to its CPM values across the cohort. For UMRs with a bimodal distribution, the density minimum between the two modes was used as the cutoff, and a call was kept only if more than 5% of samples fell on each side. SNPs were identified from the mapped UMR-seq reads. Reads were deduplicated with samtools and SNPs were called jointly for all lines against the IWGSC Chinese Spring RefSeq v1.0 genome using BCFtools ^96^ v1.15.1 (mpileup with minimum mapping quality 10 and indels skipped; call -m with diploid ploidy).

### Markers and population structure

SNPs called from UMR-seq reads (SNP), a 15K SNP-array and UMR presence/absence calls (UMR) were filtered to the same thresholds (≤ 10% missing calls; minor allele frequency > 0.05), retaining 43,622 SNPs, 10,352 SNP-array markers and 139,195 UMRs. Relationship matrices were built from each marker set^97^, compared by correlation of their off-diagonal elements and summarised by principal coordinates and Ward clustering^98^. Full details are included in **Supplementary Methods**.

### Genomic prediction

The UMR marker matrix was orthogonalised against the leading 16 SNP principal components to remove population structure shared with the SNP data. SNP-only (GBLUP) and combined SNP-plus-UMR models were fitted in ASReml-R and compared by leave-one-population-out cross-validation across five genetic groups, using predictive ability, root-mean-square error and regression bias.

### Genome-wide association

SNPs and UMRs were tested separately with the BLINK model in GAPIT^99^, with four principal components as covariates and significance at Benjamini–Hochberg FDR q ≤ 0.03. Significant UMR and SNP associations for the same trait were classified as shared when in linkage disequilibrium (r² > 0.2), and the overlap expected by chance was estimated by permutation.

## Data Availability

Sequencing reads are available through the NCBI SRA accession number xxx. Hypomethylomes can be browsed online at https://crisplab.genome.edu.au/apollo.

## ACKNOWLEDGEMENTS

The authors would like to acknowledge The University of Queensland Research Infrastructure for funding this work in the form of a Genome Innovation Hub Collaborative Project. The authors would like to acknowledge members of Genome Innovation Hub for their support, particularly Brooke Purdue and Jun Ma. We wish to acknowledge the University of Queensland’s Research Computing Centre (RCC) for its support in this research. We thank Tobin Florio for figure preparation (http://www.flozbox-science.com). We thank Nathan Springer, Steve Eichten, Milos Tanurdžić and David Kainer for invaluable discussions and feedback.

## Ethics declarations

The authors declare no competing interests.

