## Supplementary Methods and Figures for "High-throughput sequencing of unmethylated DNA (UMR-seq) reveals extensive epigenetic variation in the hypomethylome of diverse plant species"

**Illumina library preparation for UMR-seq.** Sequencing libraries were prepared using the Nextera XT DNA Library Prep Kit (Illumina, 2019). For tagmentation, 1 ng of purified DNA was used, while the volumes of all other reaction components were reduced to half those specified in the standard Nextera XT protocol. Tagmentation was performed at 55 °C for 5 min, followed by neutralization at room temperature for 5 min. Tagmented DNA was subsequently amplified in a total reaction volume of 32.5 µl, comprising 15 µl of tagmented DNA, 5 µl of each adapter and 7.5 µl of Nextera PCR Master Mix. PCR amplification was performed using the standard Nextera XT thermocycling conditions (Illumina, 2019) for 12 cycles. Following amplification, libraries were purified using SPRIselect beads according to the Nextera XT protocol, with 45 µl of beads added to 27.5 µl of amplified library. Purified libraries were quantified using the Qubit 1x High Sensitivity assay. All indexed samples were then pooled, per batch, such that each sample contributed an equal mass to the final library pool. Final library pools were sequenced on an Illumina NovaSeq S4 platform at the Australian Genome Research Facility (AGRF) using 150-bp paired-end sequencing.

**EM-seq library preparation.** For EM-seq, DNA samples were sheared (500 ng DNA in 130 uL of water) using the M220 Covaris instrument (Setting: target size-400 bp, peak incident power-140 W, duty factor-10%, 200 cycles per burst, treatment time-55 seconds, water level - 6, and water temperature - 7°C). Libraries were prepared and dual-indexed using the NEBNext Enzymatic Methyl-seq Kit (E7120S) for use with standard insert libraries (370-420 bp). The Indexing PCR was run for 6-8 cycles, cleaned and then quantified using a Qubit 4 Fluorometer. Libraries were pooled equimolar, cleaned once more and eluted in NEB elution buffer. Samples were then sequenced on an Illumina NovaSeq S4 platform at the Australian Genome Research Facility (AGRF) using 150-bp paired-end sequencing.

***In silico* digestion modelling.** *In silico* digestion modeling of the genome was performed to identify regions that would be recalcitrant to digestion by the methylation dependent enzymes due to sparse occurrences of the recognition motif. We first translated the redundant recognition motif of each restriction enzyme ($^{m}$CNNR (R = G or A) for MspJI and C$^{m}$C for FspEI) into machine-readable patterns. We then used a custom C++ script called *Grepbed* from the *seqhax* library (https://github.com/kdm9/seqhax) to find the locations of all instances of each motif in the genome, assuming every cytosine in the genome is methylated for the purpose of assessing if it could be digested. Using the known cutting patterns of each enzyme, the sites where the genome is cut are predicted and recorded, producing a table of genome fragments. The start, end and size of each fragment is then calculated. The fragments are then filtered before further analysis in two steps. In the first step, all fragments less than 100bp long were excluded, because fragments less than 100bp are size selected against during the UMR-seq library preparation. In the second step, all fragments where more than 50% of the bases are "N" are excluded. Fragments composed of N's can flag gaps in the genome assembly between contigs, so are not relevant to assessing regions of the genome recalcitrant digestion. Tandem Repeats Finder v4.09.1 [100] was then used to analyse fragments with repetitive sequences using recommended parameters. The output of the analysis is a bed file of blacklisted regions to be excluded from UMR output due to the inability to distinguish methylated DNA from unmethylated DNA using the enzymes in this protocol.

**Chromosome splitting**. The UMR-seq pipeline requires using chromosomes less than 500 Mbp long. To reduce the length of barley and wheat chromosomes, the middle-point coordinate was calculated using *seqtk* comp function [101], and a scaffold break was searched for up to 1 Mbp downstream the middle point where the split is to be done. The *seqkit* sliding function [102] was used to split chromosomes, which were then concatenated into a single genome fasta file.

**Analysis of UMR-seq and MethylC-seq Data.** UMRs identified from methylC-seq data from nine plant species were compared to normalised UMR-seq read coverage using deepTools v3.5.1 computeMatrix scale-regions (-b 2000 -a 2000, -p 2, -bs 100, --regionBodyLength 1000, --missingDataAsZero, --sortRegions descend); deepTools v3.5.1 plotHeatmap (--interpolationMethod nearest) [85].

**Analysis of UMR-seq and conservatory regions.** The UMR peak coordinates were compared with the conserved non-coding sequence (CNS) regions identified by 54), which were based on B73 version 5. Therefore, the UMR peak coordinates were lifted over to version 5 using *liftOver* (version 482) [103] and the corresponding B73V4 to V5 (Zmays_833_Zm-B73-REFERENCE-NAM-5.0) chain file, obtained from MaizeGDB (maizegdb.org). BEDtools *intersect* was then used to quantify the number of both UMR peaks overlapping the CNS regions and, conversely, CNS regions overlapping the UMR peaks [92]. Additionally, *regioneR* v1.44.0 was used to assess the enrichment of CNS regions within UMRs by comparing the observed number of overlaps with the expected distribution generated from 50 permutations [93]. The UMR-seq:CNS heatmap was produced with deepTools as described above. To visualize CNS-associated UMR-seq signal, normalised UMR-seq read coverage tracks were processed using deepTools v3.5.5 *computeMatrix* (scale regions -b 2000 -a 2000, -p 8, -bs 50, –missingDataZero, –sortRegions descend). The resulting signal metrics were visualized as heatmaps using deepTools v3.5.5 *plotHeatmap* (--interpolationMethod bilinear) [85].

**Analysis of UMR-seq and TF binding sites.** The overlap analysis, and permutation-testing analysis workflow described above for CNS regions was applied to the transcription factor (TF) binding sites. Briefly, BEDTools *intersect* was used to quantify overlaps between UMRs and TF binding sites and, *regioneR* v1.44.0 was used to assess enrichment by comparing the observed overlap with the expected distribution generated from 5,000 permutations [92,93].

**Population UMR analysis in wheat and the pan-hypomethylome.** Briefly, the panel comprised 191 elite European winter wheat cultivars representing almost five decades of breeding progress. Genotypes were phenotyped across six field locations over two seasons (not all traits were scored in every environment; radiation-use efficiency was measured at one location) under three management intensities: high nitrogen with high fungicide (HiN.HiF), high nitrogen without fungicide (HiN.NoF), and low nitrogen without fungicide (LoN.NoF). Nitrogen treatments were applied at a total nitrogen supply rate of 220 kg N $ha^{-1}$ (high) and 110 kg N $ha^{-1}$ (low), adjusted for soil-mineralised nitrogen, and the high-intensity treatment received fungicides, insecticides and growth regulators at standard regional practice[69]. Traits spanning yield components, development (heading date and plant height), grain quality, disease resistance, nitrogen- and radiation-use efficiency (n = 53 trait-management combinations) were phenotyped following the described methods and analysis framework[69]. Adjusted means in the form of best linear unbiased estimators (BLUEs) were taken directly, estimated for each line in each management regime across years, locations and replicates with a linear mixed model in which genotype, management and their interaction were fitted as fixed effects and year, location, their interaction, replicate and block as random effects[69] and used for all downstream analysis. Of the 191 genotypes in the original panel, 187 were included in the present study following sample availability and quality control.

**Population structure and relatedness analyses.** Three marker sets were compared for relatedness and population structure: SNPs called from the UMR-seq pipeline (SNP), an independent 15K SNP

Illumina Infinium iSelect genotyping array (SNP-array), and UMR presence/absence calls coded 0/2 (UMR). All three sets were restricted to the 187 analysed lines before filtering, and the same thresholds were applied to each: markers with > 10% missing calls and markers with MAF ≤ 0.05 (computed after imputation) were removed. Missing SNP genotypes were imputed by k-nearest-neighbour imputation (k = 10; using genome-wide allele-difference distance and neighbours weighted by inverse distance). Filtering retained 43,622 of 49,865 SNPs, 10,352 of 13,006 SNP-array markers and 139,195 of 142,391 UMRs. Marker distribution along the chromosomes (**Supplementary Fig. 19**) was summarised as the number of markers with a chromosome position per 5 Mbp window (IWGSC RefSeq v1.0; chromosome length approximated by the largest marker position; 8,781 of the 10,352 SNP-array markers had a chromosome position in the supplied map), expressed relative to each marker set's genome-wide mean. Concordance between the SNP and SNP-array distributions was measured by the Spearman correlation of window counts.

Genomic and methylation relationship matrices were constructed from each marker set using the VanRaden method[97]. Agreement between relationship matrices was summarised by the Pearson correlation of their off-diagonal elements. To reduce the population structure shared with the SNP data before genomic prediction, the UMR marker matrix was orthogonalized against the SNP principal-component space: the centred UMR marker matrix was projected off the leading k eigenvectors of the SNP relationship matrix, and a residual UMR relationship matrix was constructed from the remaining variation. k was set at the plateau of the SNP scree curve, defined as the principal component (among PCs 1–50) with the largest perpendicular distance from the straight line joining the cumulative-variance curve at PC1 and PC50, giving k = 16 (42.3% of SNP variance).

Population structure was visualised by eigendecomposition of each relationship matrix, with individuals projected onto the first two principal coordinates. Individuals were assigned to three groups by Ward's minimum-variance hierarchical clustering[98] (Ward.D2) of the leading 10 SNP principal coordinates, and this common grouping was applied across all panels for visual comparison and both relationship matrices were displayed in the order of the single SNP dendrogram. Agreement between the SNP and SNP-array relationship matrices (**Supplementary Fig. 18**) was summarised by off-diagonal Pearson correlations.

**Genomic prediction and heritability estimation.** Genomic prediction models were fitted in ASReml-R (v4.2.0.332) using best linear unbiased estimates (BLUEs) per trait–management (described above). The UMR kernel was the orthogonalised UMR relationship matrix described above and the SNP-derived relationship matrix was used as the genomic kernel. For comparability, each kernel was scaled to an average semivariance of one. For each of the 53 trait–management combinations, two models were fitted: a SNP-only (standard GBLUP) model and a combined multi-kernel model. The combined multi-kernel model was specified as:

$$y = \mu + Z_{gSNP} + Z_{gUMR} + e$$

where y is the vector of trait BLUEs, μ the overall mean, and Z the incidence matrix linking observations to individuals. The SNP and methylation genetic effects, $_{gSNP}$ and $_{gUMR}$, were fitted as independent random effects with covariance structures $G\sigma^2_1$ and $K\ \sigma^2_2$ respectively, where G is the SNP-derived relationship matrix, K the orthorthogonalised UMR kernel, and $\sigma^2_1$ and $\sigma^2_2$ their associated variance components. The residuals were distributed as $I\sigma^2_e$. The GRM-only model was identical but excluded the term. Narrow-sense heritability of each trait–management combination was estimated from the SNP-only model as $h^2 = \sigma^2_g / (\sigma^2_g + \sigma^2_e)$, where $\sigma^2_g$ is the SNP genetic variance and $\sigma^2_e$ the residual variance.

Predictive performance was evaluated by cross-validation across population groups, comparing the SNP-only model with the combined multi-kernel model. Individuals were partitioned into five groups (31–49 lines) by partitioning around medoids (PAM) clustering of SNP-derived genetic dissimilarity (one minus the correlation matrix of the SNP relationship matrix), and each group was masked and predicted from the other four in turn (leave-one-population-out). For each fold, four metrics were calculated: predictive ability (r) calculated as the Pearson correlation between predicted breeding values and observed phenotypes, prediction accuracy as ($r_{PA}$) divided by the square root of the SNP-only model heritability, root-mean-square error, and regression bias as the slope of the regression of observed on predicted values.

**Genome-wide association analyses.** Separate genome-wide association analyses were performed for the SNP and UMR marker sets using GAPIT (v3) in R, and datasets formatted according to documentation specifications [99]. Associations were tested for each of the 53 trait-management combinations, using BLUEs as the response. Only markers with a chromosome position were tested (42,905 SNPs and 132,433 UMRs). To account for population structure, the first four principal components (PCs) of each marker set were fitted as covariates. Beyond PC4, each additional component explained ≤ 1.92% of UMR variance and ≤ 2.82% of SNP variance. Association testing used the BLINK model. Per-marker P-values were corrected for multiple testing within each trait × marker-set analysis using the Benjamini–Hochberg false discovery rate (FDR), and associations were declared significant at an FDR-adjusted $q \leq 0.03$. For each trait, a significant UMR and a significant SNP on the same chromosome were classified as shared if their genotypes were in linkage disequilibrium (LD; $r^2 > 0.2$); linked associations were grouped into loci, and UMR associations not linked to any significant SNP were counted as not in LD with a SNP association. The number of overlaps expected by chance was estimated by replacing each significant UMR (or, separately, each significant SNP) with a randomly drawn tested marker of the same type on the same chromosome and in the same MAF quintile (1,000 permutations), giving the expected count, its 95% interval and a one-sided permutation P. To assess SNP coverage around significant UMRs, the number of tested SNPs within 1 Mbp and the maximum $r^2$ with any tested SNP within 1 Mbp were compared between the 163 distinct significant UMRs and 1,000 sets of tested UMRs matched by chromosome and MAF quintile; maximum $r^2$ is summarised for UMRs with at least one SNP within 1 Mbp. Association counts were summarised by trait class (**Supplementary Table 7**).

### SUPPLEMENTARY TABLES

Table and legends attached.

### SUPPLEMENTARY FIGURES

Figures and legends below.

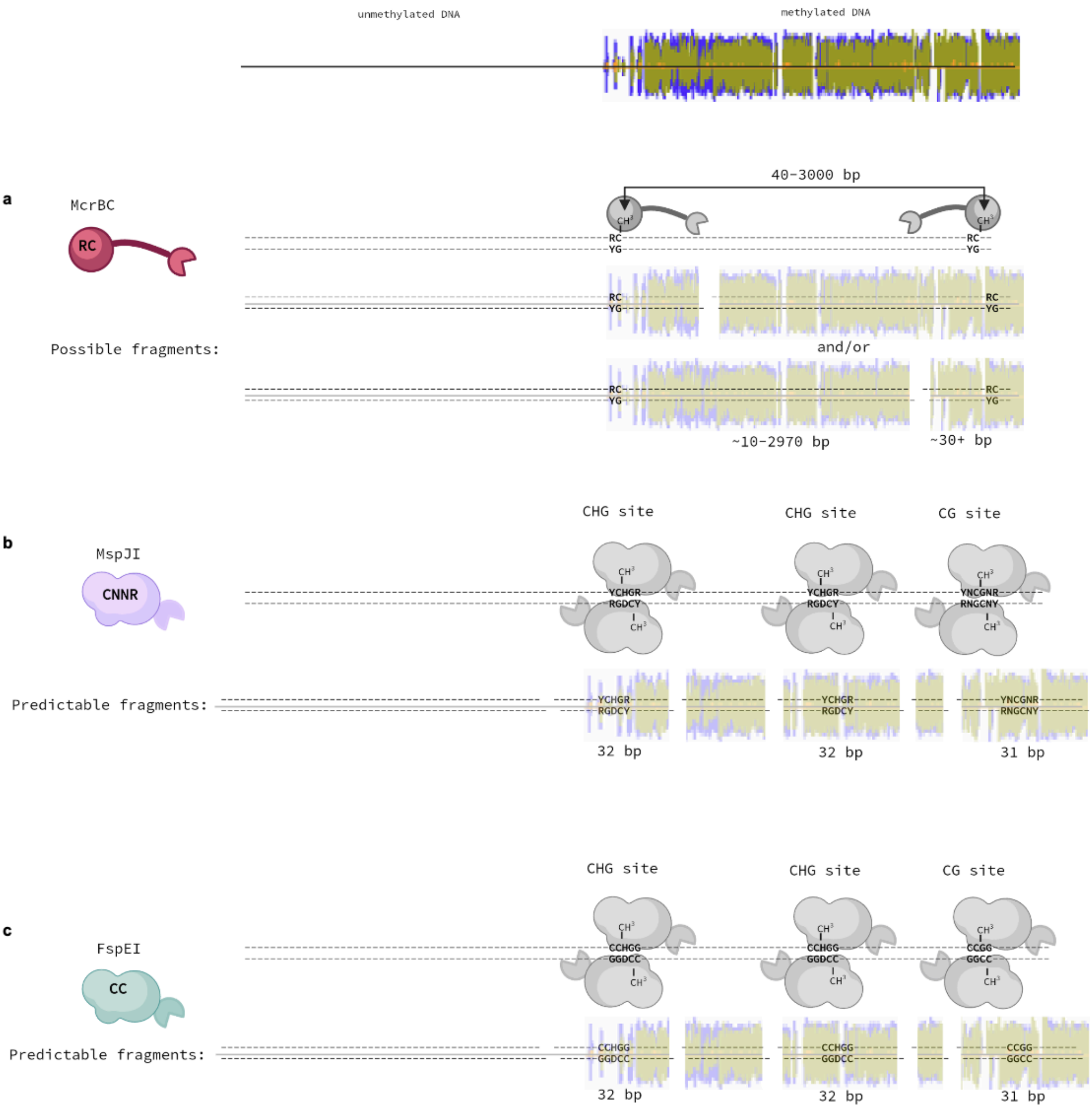


**Supplementary Figure 1. DNA methylation dependent restriction enzymes, recognition sequences and digest fragments. a**, McrBC enzyme complex originating from *Escherichia coli* (Ec) is formed by a EcMcrB hexamer and EcMcrC which together recognise RmC sites (where R is a purine) that can be fully or hemi-methylated and catalyses cleavage. McrBC needs at least two RmC sites that can be 40-3000 base pairs apart, the complex cleaves one or both DNA strands at approximately 30bp away from either site. **b**, The MspJI restriction enzyme recognises 5mCHG and 5mCG sites, where sites are symmetrically methylated. Cleavage of DNA occurs downstream of both sites with a cut made 12bp downstream on the top strand and 16bp away on the bottom strand, a 31 to 32 bp fragment is generated from this enzyme. **c**, FspEI behaves similarly to MspJI, FspEI recognises symmetrical CmC sites which encompasses CG methylation and CHG methylation. Methylation can be either C5-methylation (5mC) or C5-hydroxymethylation (5hmC). Cleavage occurs downstream from each site at 12bp and 16pb for both DNA strands, creating a 31-32 bp fragment. Created in BioRender.

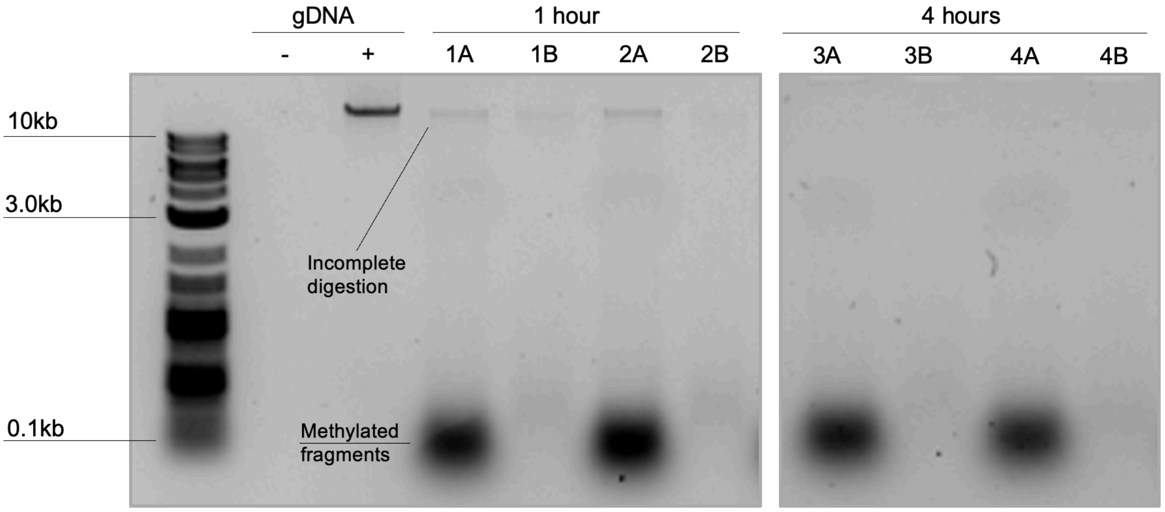


**Supplementary Figure 2. Gel electrophoresis of digested DNA comparing optimally digested samples (4 hr) to under-digested samples (1 hr) each visualised before and after size selection.** Genomic DNA (left) is digested by modification-dependent endonucleases MspJI and FspEI resulting in cleavage of methylated DNA into 31-32bp bp fragments and resulting in a range of larger unmethylated DNA fragments appearing as a smear (lanes 1A, 2A, 3A, 4A). Paramagnetic beads (bead:DNA ratio 0.7x) were used to isolate the higher molecular weight unmethylated DNA from the low molecular weight methylated fraction (lane 1B, 2B, 3B, 4B) thereby removing methylated DNA from the sample. A high concentration of fragments smaller than 100 bp are visible in lanes labelled "A", which are removed by bead-based size selection shown in "B". Complete digestion of gDNA is shown among samples 3 and 4 digested for 4 hours in comparison to samples 1 and 2, digested for 1 hour with evidence of a high molecular weight genomic DNA band indicative of incomplete digestion of gDNA. Optimal digestion time can depend on the fraction of UMRs in a genome, which is species dependent, sample characteristics such as starting mass and sample quality, and contaminants may alter enzyme activity. This sample is from barley, we found that the optimal digest time for the samples presented in the work was 120 minutes, but provide this as an example of under and complete digestion. Digests should be examined (and adjusted if needed) for each experiment.

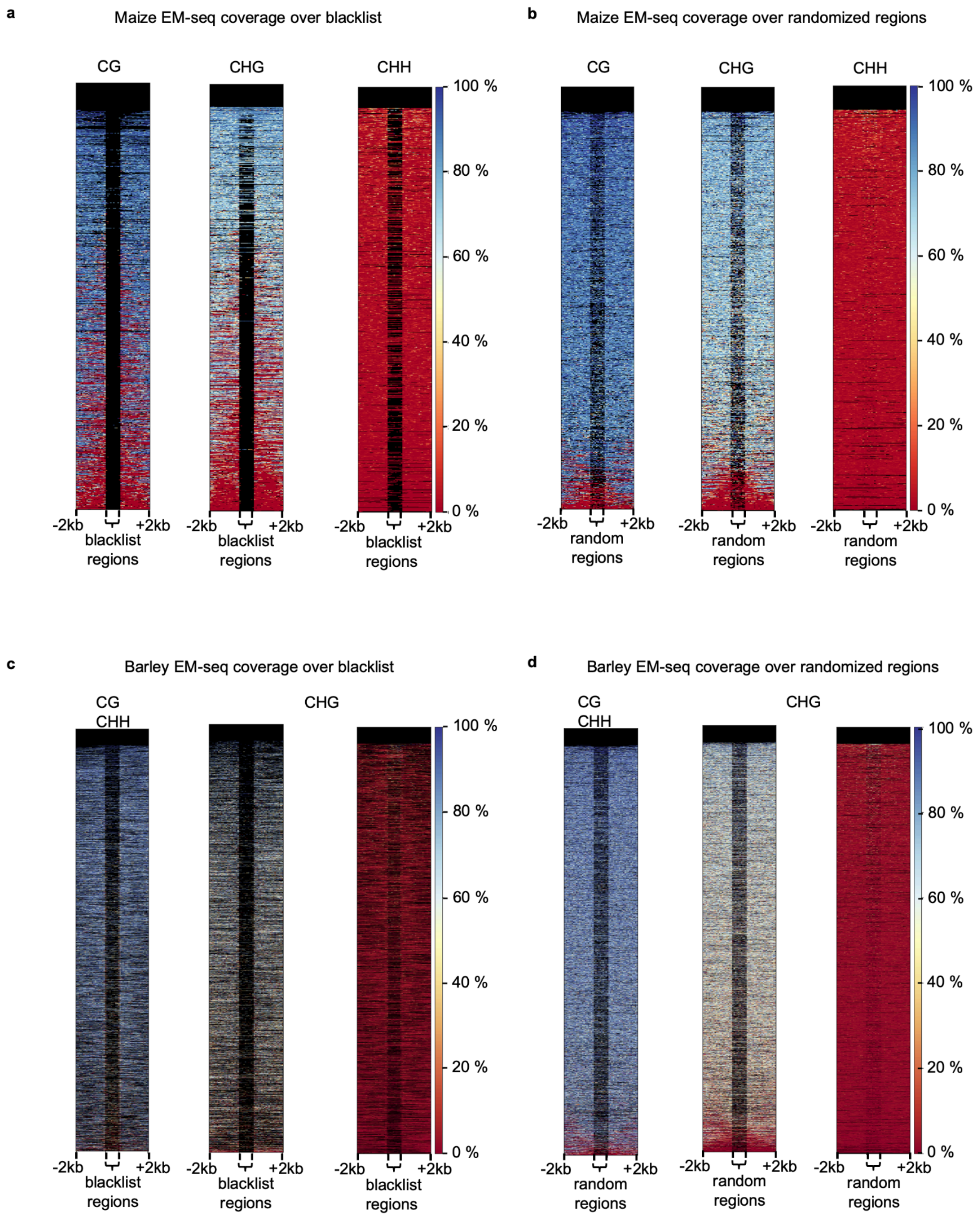


**Supplementary Figure 3. DNA methylation levels determined by EM-seq at regions recalcitrant to UMR-seq digestion. a**, Maize (genotype B73) EM-seq data for CG, CHG and CHH methylation contexts at blacklisted recalcitrant regions identified from the *in silico* digest of the maize reference genome. Most regions lack EM-seq data (black), indicating they are recalcitrant to both UMR-seq and EM-seq analysis. **b**, Maize methylation in all contexts over regions randomly selected (but equivalent in size and number to the recalcitrant regions) as a null control for expected pattern if regions are randomly chosen compared to the blacklist. Random regions have higher average methylation than recalcitrant regions. **c**, Barley (genotype Morex) EM-seq data for

CG, CHG and CHH methylation contexts at blacklisted regions identified from the *in silico* digest of the barley Morex reference genome. **d**, Barley EM-seq data over randomly generated regions. Blue indicates high levels of DNA methylation, red low levels and black indicates missing data mostly due to low mapability of repetitive regions.

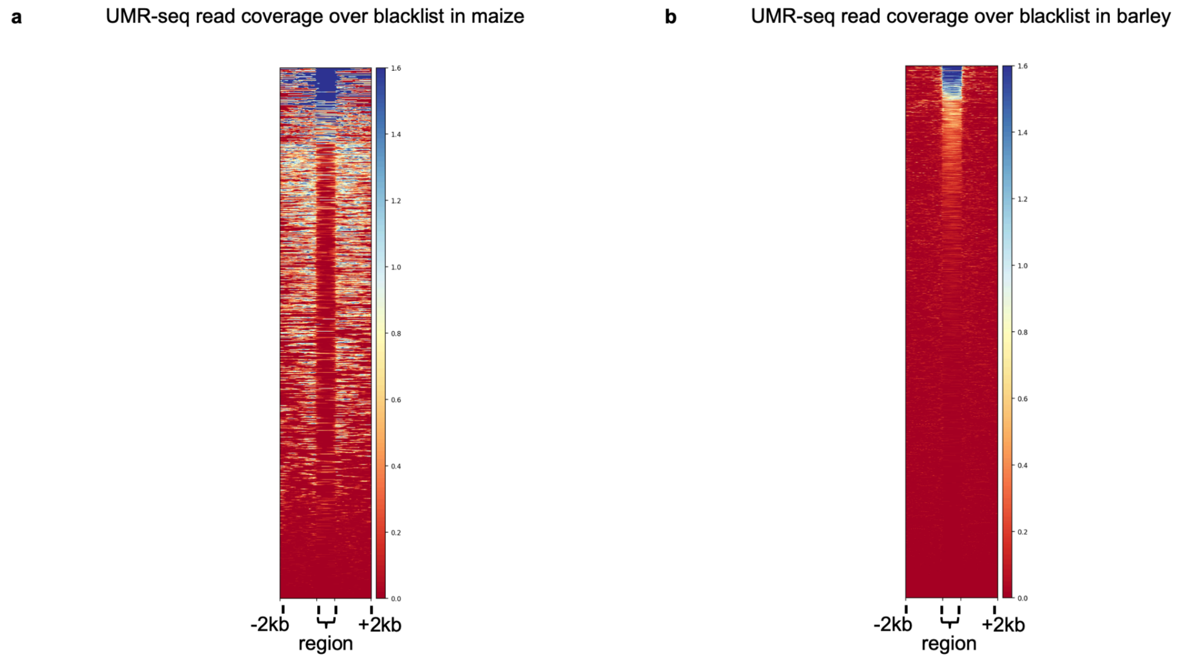


**Supplementary Figure 4. UMR-seq read coverage at regions identified as recalcitrant to UMR-seq digestion. a**, UMR-seq read coverage is visualised over regions in the blacklist containing fragments recalcitrant to digestion with MspJI and FspEI for maize (genotype B73). Most regions lack data (red), although a small number have some read coverage (blue). **b**, UMR-seq read coverage over the blacklist generated for barley (genotype Morex). Read coverage is visualised over the blacklisted region and 2000 bp either side, blue indicates high read coverage (CPM) and red indicates no reads.

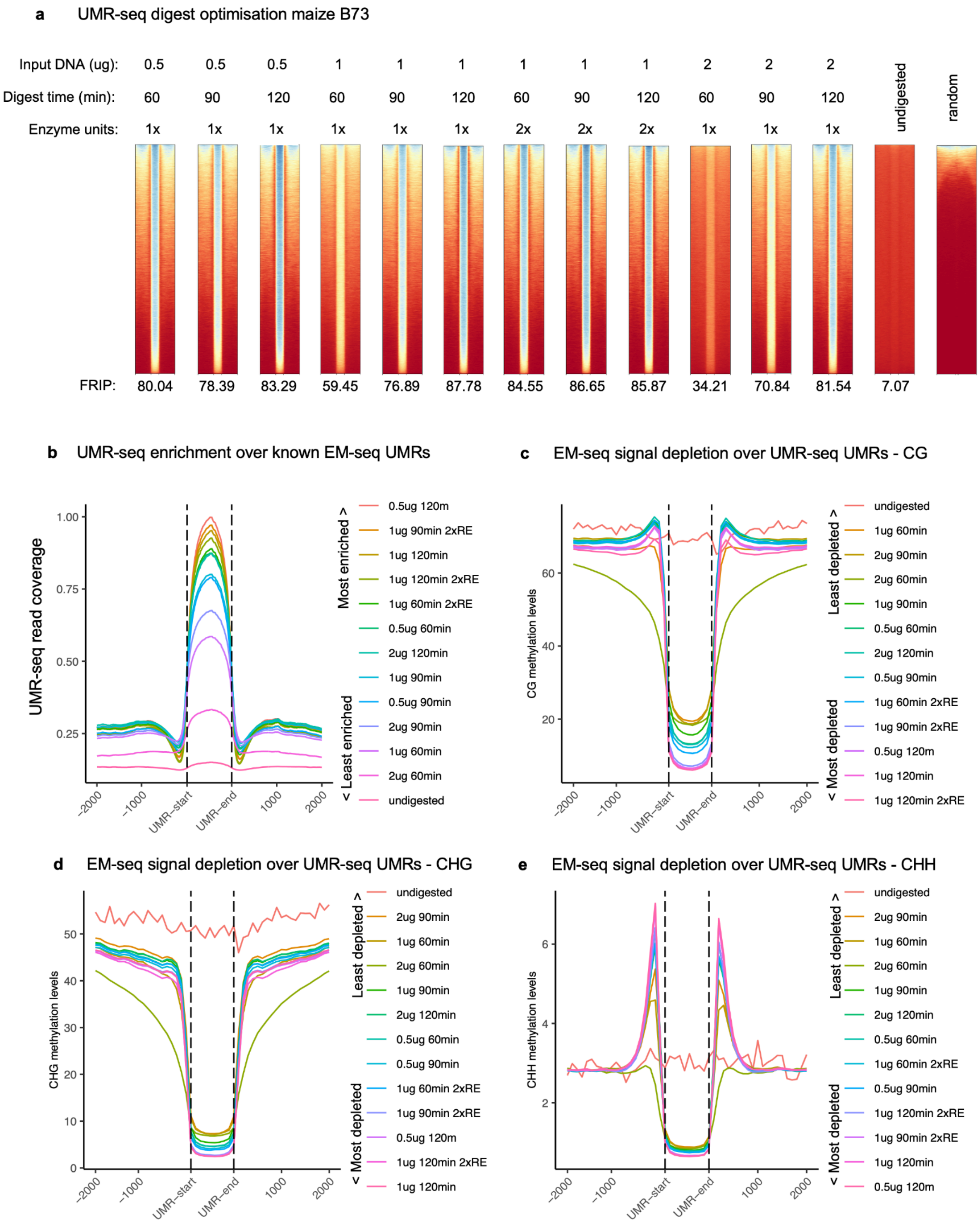


**Supplementary Figure 5. Digestion optimisation of genomic DNA from maize (cv. B73).** UMR-seq is robust over a range of digest conditions, however, we observed clear under digestion when DNA input was too high and digest time (or enzyme units) too low. We did not observe an impact of overdigestion on library quality. Overdigestion may occur if digestion time is extended further. We did observe that extended digestion can result in low yield of DNA post digestion, indirectly impacting the ability to construct a quality library with sufficient complexity.

a) The enzyme digest conditions for UMR isolation were investigated by varying the mass of input DNA, digest time and enzyme units. UMR-seq read coverage was then visualised over known UMRs identified using EM-seq. 1x enzyme units refers to 5 units. FRIP calculated using *plotEnrichment* function in the *deeptools* package by determining the fraction of reads in peaks (identified using epic2, 50 bp sliding window). The undigested profile is a negative/input control, prepared by omitting the enzymes from the digestion but otherwise preparing a sequencing library in the same way as UMR-seq libraries are prepared (so essentially this sample is a Nextera whole genome sequencing library prepared from gDNA). Random regions sampled from the genome using bedtool shuffle using the methylC-seq UMR bed file as input. b) Metaplot of UMR-seq read coverage over known UMRs. c) CG context DNA methylation levels determined using EM-seq data for UMR-seq UMRs. MACS2 was for peak calling (narrow peak mode) to identify UMR-seq UMRs owing to its higher stringency excluding partially methylated regions (see Supp Figure 6). Sample key is ordered by level of enrichment. d) CHG context DNA methylation metaplot and e) CHH context metaplot.

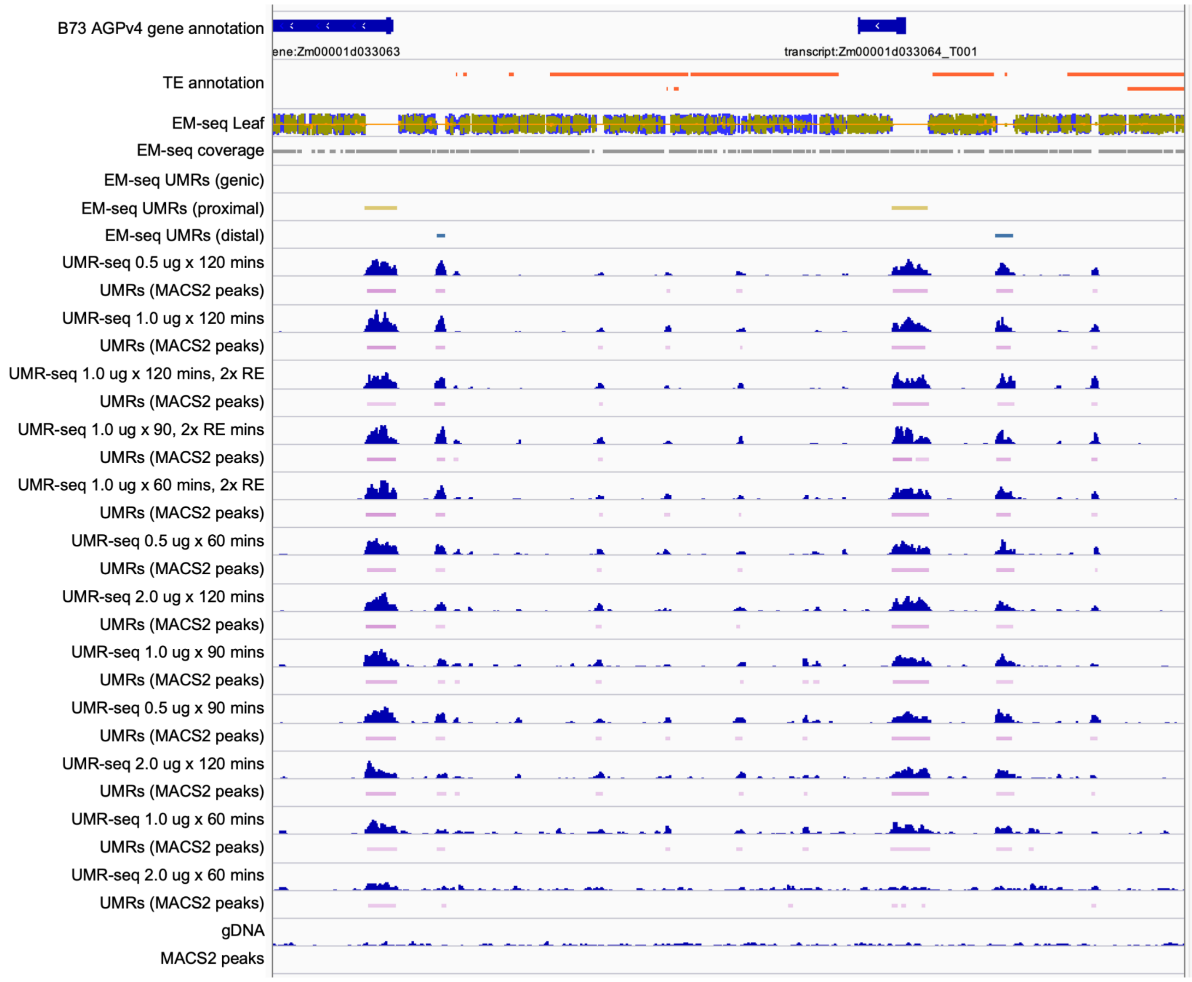


**Supplementary Figure 6. Browser example of digestion optimisation in maize B73.** Example UMR-seq profiles for the digestion optimisation in maize. UMR-seq read coverage shown in dark blue (reads normalised to CPM); UMRs identified using MACS2 depicted as pink bars below the coverage tracks. Samples ordered by UMR-seq enrichment (library quality) based on Supp Figure 7. UMR-seq coverage is observed to be highly reproducible over a range of digest conditions; however, underdigestion is observed when DNA input is high and digest time too low, for example, 2.0 µg digested 60 minutes gave a very low quality library with insufficient enrichment for UMRs.

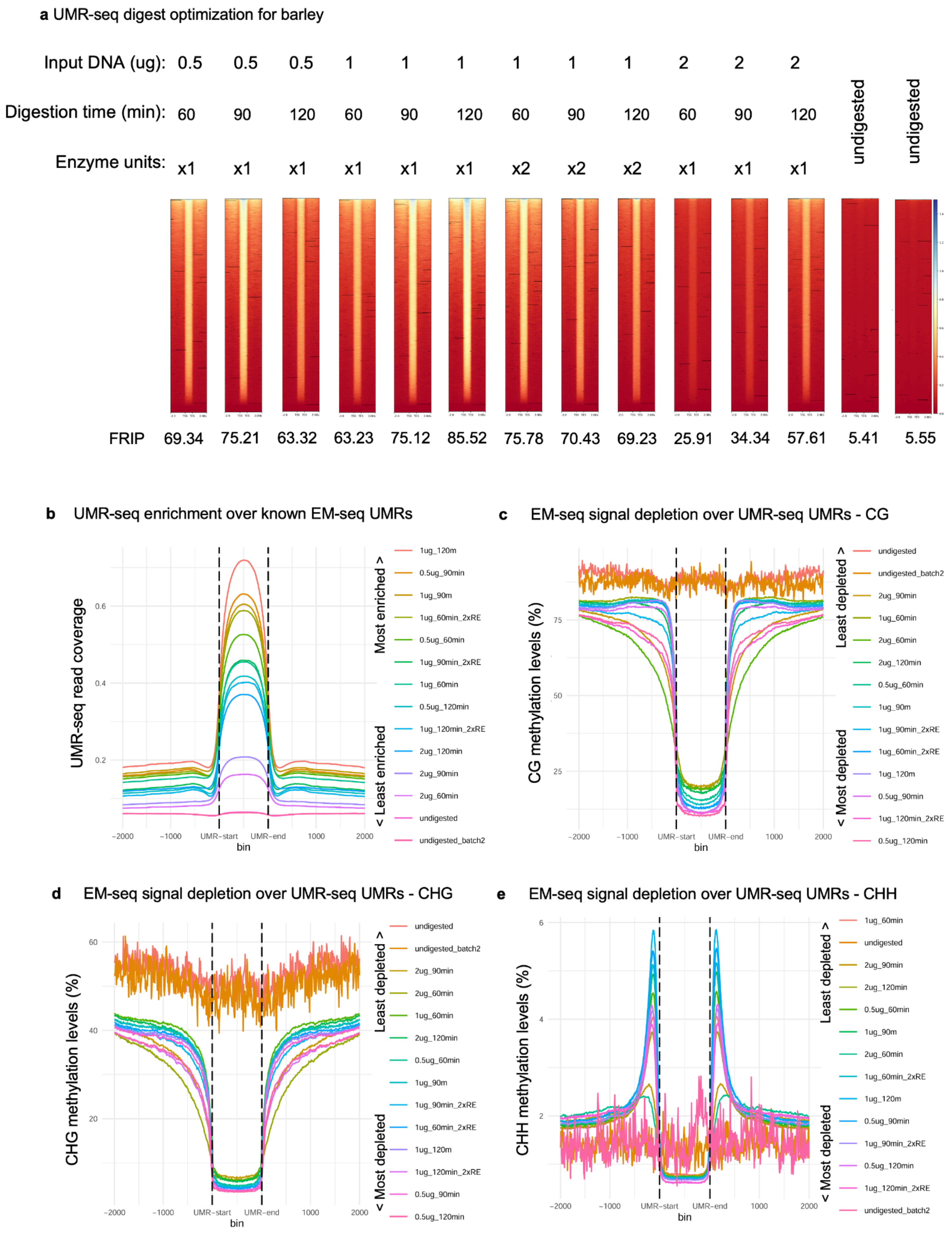


**Supplementary Figure 7. Digestion optimisation of genomic DNA from barley (cv. Golden Promise).** As in maize, we found that UMR-seq is robust over a range of digest conditions in barley. Here barley genotype Golden promise was used for digestion optimisation owing to the availability of DNA samples and the desire to optimise UMR-seq in a highly transformable variety for future functional analyses. Golden promise UMR-seq was compared to Morex EM-seq data because we had available a much higher quality EM-seq data set for Morex compared to a low quality EM-seq data for Golden promise. We observed clear under digestion when

DNA input was too high and digest time (or enzyme units) too low, for example 2 µg digested for 60 or 90 minutes. Again we did not observe an obvious impact of overdigestion on library quality. We did observe that extended digestion can result in low yield of DNA post digestion, which was more of an issue in barley owing to the smaller fraction of the genome (gDNA) that is unmethylated compared to maize. **a,** The enzyme digest conditions for UMR isolation were investigated by varying the mass of input DNA, digest time and enzyme units. UMR-seq read coverage was then visualised over known UMRs identified using EM-seq. 1x enzyme units refers to 5 units. FRIP calculated using *plotEnrichment* function in the *deeptools* package by determining the fraction of reads in peaks (identified using epic2, 50 bp sliding window). The undigested profile is a negative/input control, prepared by omitting the enzymes from the digestion but otherwise preparing a sequencing library in the same way as UMR-seq libraries are prepared (so this sample is a Nextera whole genome sequencing library prepared from gDNA). Random regions sampled from the genome using bedtool shuffle using the methylC-seq UMR bed file as input. **b,** Metaplot of UMR-seq read coverage over known UMRs. **c,** CG context DNA methylation levels determined using EM-seq data for UMR-seq UMRs. MACS2 was used for peak calling (narrow peak mode) to identify UMR-seq UMRs owing to its higher stringency excluding partially methylated regions (see Supplementary Figure 12). Sample key is ordered by level of enrichment. **d,** CHG context DNA methylation metaplot and **e,** CHH context metaplot.

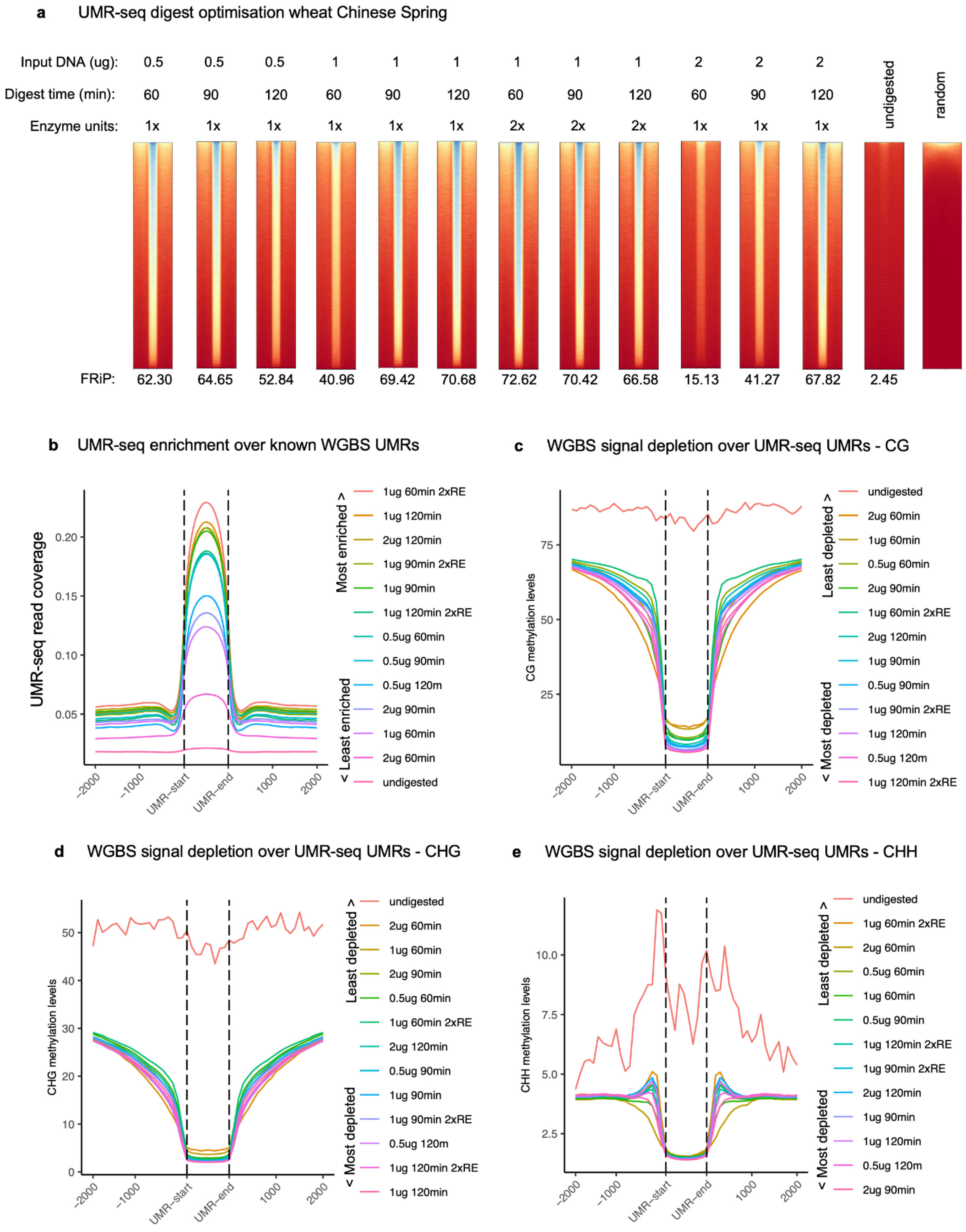


**Supplementary Figure 8. Digestion optimisation of genomic DNA from wheat (cv. Chinese Spring).** As in maize and barley, we found that UMR-seq is robust over a range of digest conditions in wheat. We observed clear under digestion when DNA input was too high and digest time (or enzyme units) too low. Again we did not observe an obvious impact of overdigestion on library quality. We did observe that extended digestion can result in low yield of DNA post digestion, which was more of an issue in wheat owing to the smaller fraction of the genome (gDNA) that is unmethylated compared to maize. **a**, The enzyme digest conditions for UMR isolation were investigated by varying the mass of input DNA, digest time and enzyme units. UMR-seq read coverage was then visualised over known UMRs identified using published WGBS. 1x enzyme units refers to 5 units. FRiP calculated using *plotEnrichment* function in the *deeptools* package by determining the fraction of reads in peaks (identified using epic2, 50 bp sliding window). The undigested profile is a negative/input control, prepared by omitting the enzymes from the digestion but otherwise preparing a sequencing library in the same way as UMR-seq libraries are prepared (so this sample is a Nextera whole genome sequencing library prepared from gDNA). Random regions sampled from the genome using bedtools shuffle using the methylC-seq UMR bed file as input. **b**, Metaplot of UMR-seq read coverage over known UMRs. **c**, CG context DNA methylation levels determined using WGBS data for UMR-seq UMRs. MACS2 was used for peak calling (narrow peak mode) to identify UMR-seq UMRs owing to its higher stringency, excluding partially methylated regions (see Supp Figure 12). Sample key is ordered by level of enrichment. **d**, CHG context DNA methylation metaplot and (**e**) CHH context metaplot.

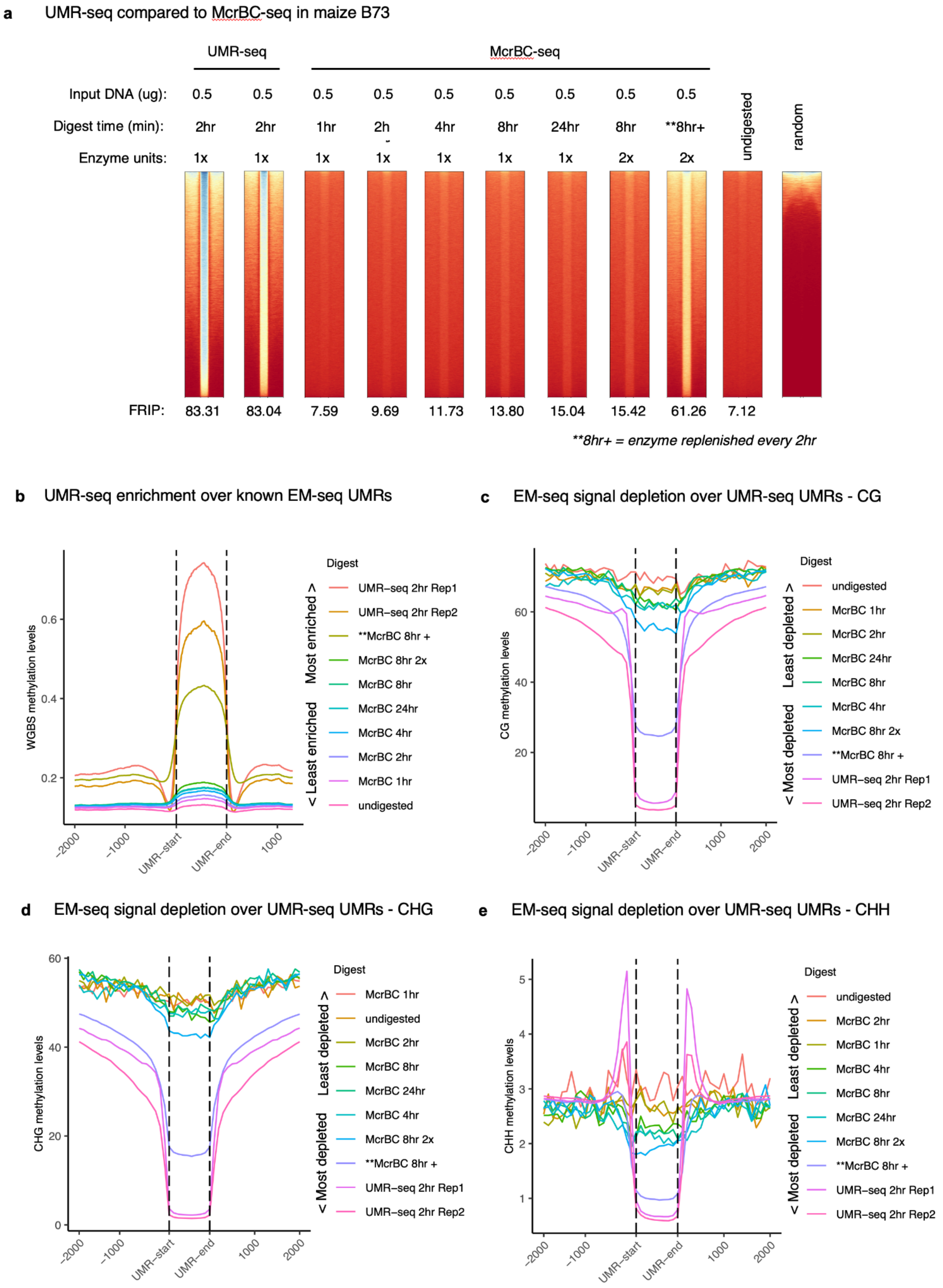


**Supplementary Figure 9. UMR-seq compared to McrBC-seq.** McrBC-seq libraries for maize B73 were prepared in the same manner as UMR-sqe libraries, except McrBC was used for methylated DNA digestion in place of FspEI and MspJI. Performance of McrBC was evaluated and compared to UMR-seq by determined read coverage over known EM-seq UMRs, visualised in **a,** heatmaps annotated with FRIP scores, and **b,** as metaplots. **c,** CG, **d,** CHG and **e,** CHH context DNA methylation levels determined using EM-seq data for mcrBC-seq UMRs (MACS2). Sample key is ordered by level of enrichment. McrBC exhibited the best enrichment for UMRs and reasonable depletion of methylated DNA when the digest is performed over 24 hrs with fresh enzyme added every 8 hrs. However, UMR-seq had better performance and only required a 2-hour digestion. Shorter McrBC digestions showed relatively poor depletion of methylated DNA.

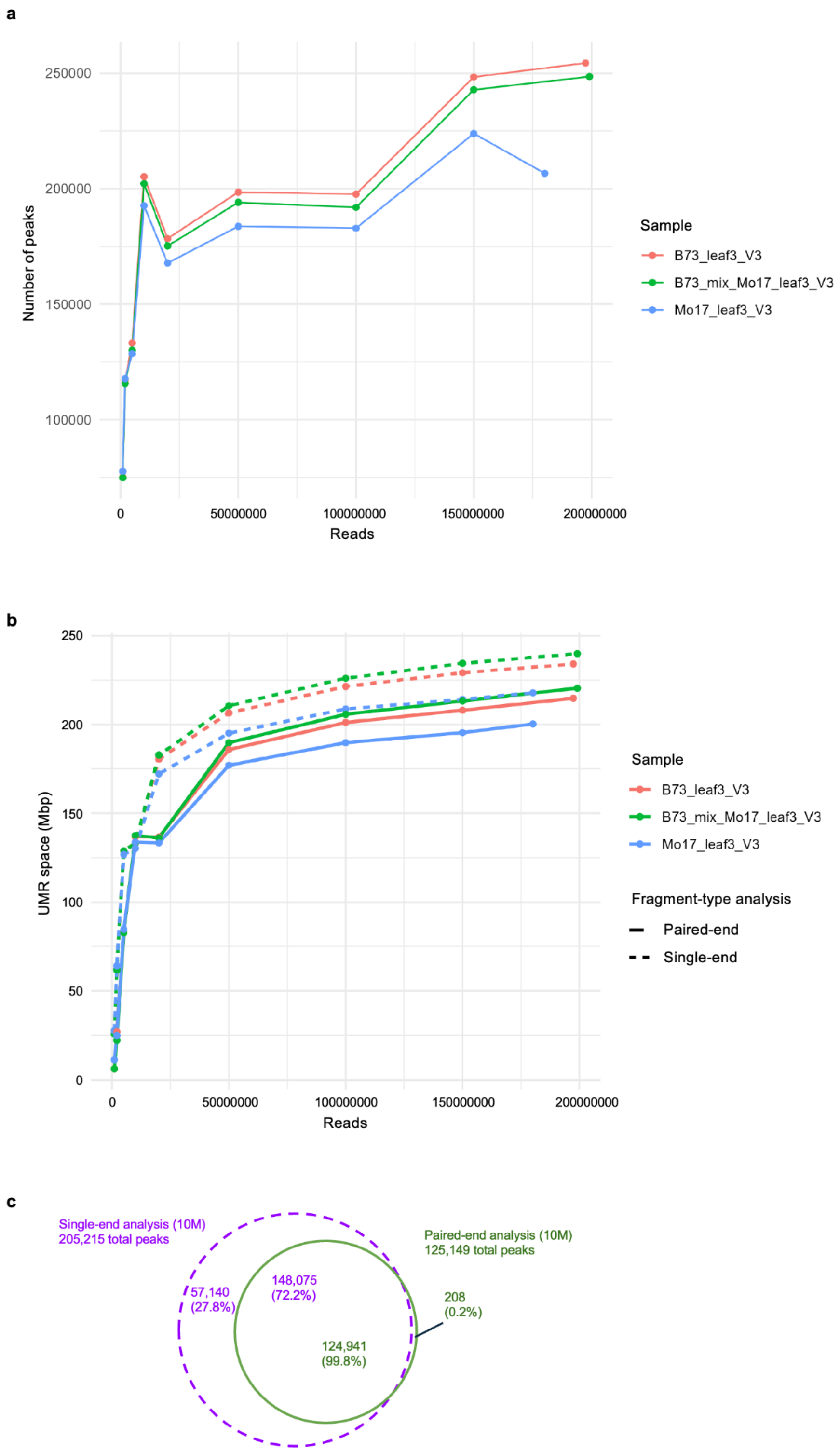


**Supplementary Figure 10. Analysis of the effect of sample read counts in UMR peak calling and UMR space, in subsets of deep-sequencing datasets from maize leaf samples. a,** The number of detected peaks increases rapidly with read count up to 10 million reads, then plateaus until 100 million reads and increases again at 150 million reads. **b**, UMR space total size, in megabase pairs, increases sharply with total read number until 20 million reads, after which the UMR space gains are reduced. We also noticed that the paired-end mode of epic2, introduced in a recent update, detects fewer peaks than the single-end analysis function in all samples, at the same read count. **c**, Representative Venn diagram showing that the peaks detected using the single-end analysis function of epic2 (in purple) encompass 99.8% of the peaks called using the paired-end mode (in green). The samples are from two genotypes of maize, Mo17 and B73, and an artificial heterozygote created by mixing genomic DNA from B73 and Mo17 in a 50:50 ratio, demonstrating the consistency of the analysis across samples.

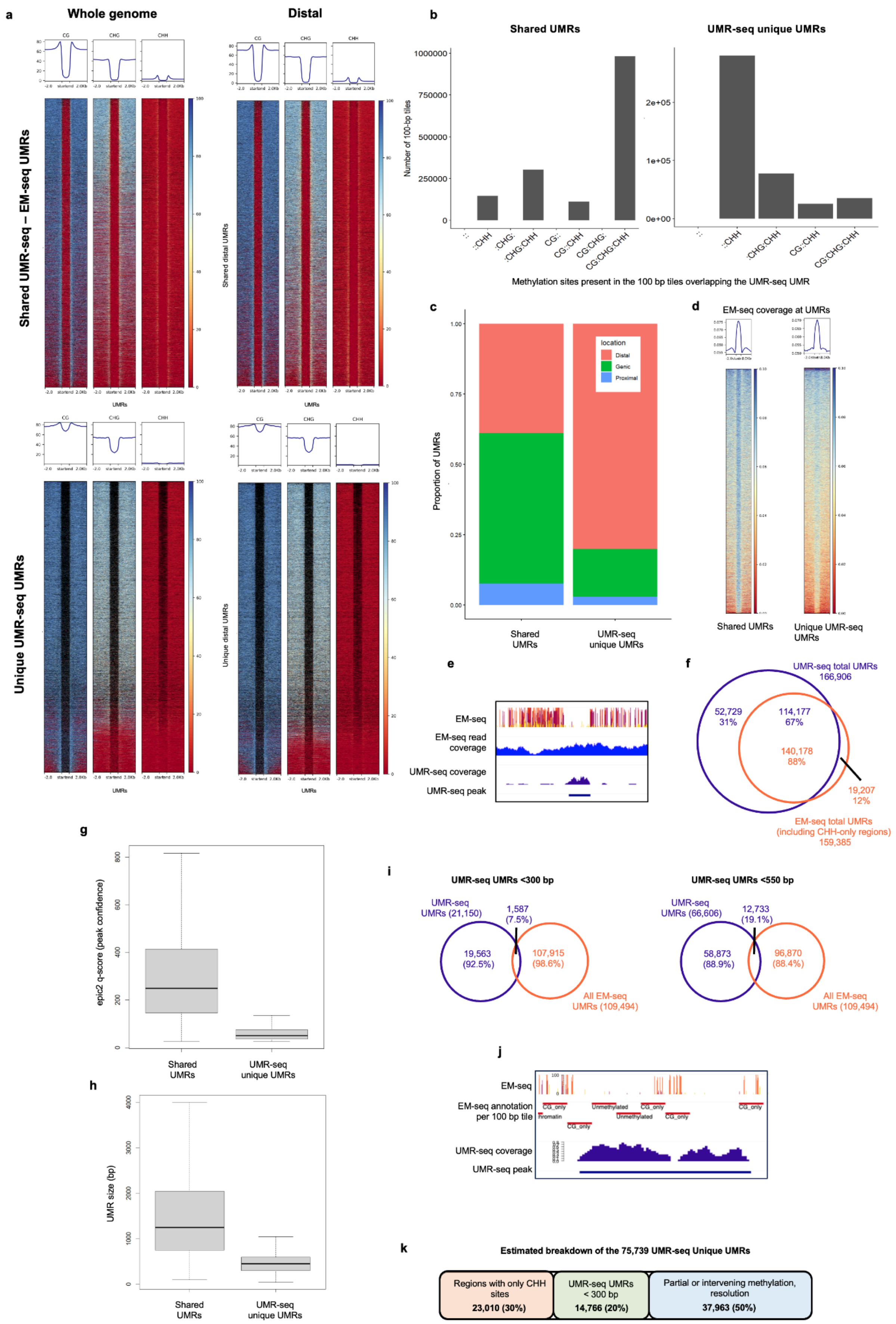

a
Whole genome
Distal
CG
CHG
CHH
Shared UMR-seq – EM-seq UMRs
Shared distal UMRs
Unique UMR-seq UMRs
Unique distal UMRs
UMRs
b
Shared UMRs
UMR-seq unique UMRs
Number of 100-bp tiles
1000000
750000
500000
250000
0
2e+05
1e+05
0e+00
::
::CHH
:CHG:
:CHG:CHH
CG::
CG::CHH
CG:CHG:
CG:CHG:CHH
Methylation sites present in the 100 bp tiles overlapping the UMR-seq UMR
c
Proportion of UMRs
1.00
0.75
0.50
0.25
0.00
location
Distal
Genic
Proximal
Shared UMRs
UMR-seq unique UMRs
d
EM-seq coverage at UMRs
Shared UMRs
Unique UMR-seq UMRs
e
EM-seq
EM-seq read coverage
UMR-seq coverage
UMR-seq peak
f
UMR-seq total UMRs 166,906
52,729 31%
114,177 67%
140,178 88%
19,207 12%
EM-seq total UMRs (including CHH-only regions) 159,385
g
epic2 q-score (peak confidence)
800
600
400
200
0
Shared UMRs
UMR-seq unique UMRs
h
UMR size (bp)
4000
3000
2000
1000
0
Shared UMRs
UMR-seq unique UMRs
i
UMR-seq UMRs <300 bp
UMR-seq UMRs (21,150)
1,587 (7.5%)
19,563 (92.5%)
107,915 (98.6%)
All EM-seq UMRs (109,494)
UMR-seq UMRs <550 bp
UMR-seq UMRs (66,606)
12,733 (19.1%)
58,873 (88.9%)
96,870 (88.4%)
All EM-seq UMRs (109,494)
j
EM-seq
EM-seq annotation per 100 bp tile
CG_only
Unmethylated
UMR-seq coverage
UMR-seq peak
k
Estimated breakdown of the 75,739 UMR-seq Unique UMRs
Regions with only CHH sites 23,010 (30%)
UMR-seq UMRs < 300 bp 14,766 (20%)
Partial or intervening methylation, resolution 37,963 (50%)

**Supplementary Figure 11. Characterisation of UMRs identified in UMR-seq but not in EM-seq. a,** EM-seq methylation levels at each cytosine context are visualised over UMRs identified by both EM-seq and UMR-seq ("Shared UMRs") (top panels) and UMR-seq UMRs that do not overlap EM-seq UMRs ("UMR-seq unique UMRs") (bottom panels), for UMR-seq UMRs across the whole genome (left), and distal UMRs (right). In the heatmaps visualising the whole genome context, the UMR-seq UMRs occur in between regions methylated in the CG and CHG context (see peaks neighbouring the UMRs in the metaplots), these may represent small UMRs surrounded by methylated regions, due to the finer resolution of UMR-seq compared to EM-seq (100 bp resolution and filtered to UMRs 300 bp and greater). The majority of UMR-seq unique UMRs occur in regions that, when investigated in EM-seq data, have low levels of CHH methylation and some have partial/sparse methylation in the CG and CHG context but are mainly characterised by insufficient cytosine data represented by the black shading. **b**, Bar plot showing the presence of cytosines in each context in 100 bp-tiles overlapping UMR-seq UMRs: in Shared UMRs (left) cytosines are abundant in all contexts, whereas Unique UMR-seq UMRs (right) contain cytosines mostly in CHH context. **c,** The genomic distribution of UMR-seq unique UMRs is enriched in intergenic gene distal regions (genic refers to an overlap with a gene locus defined by the TSS and TTS; proximal refers to an overlap with the region 2 kbp upstream of the TSS, or 2 kbp downstream of the TTS; distal refers to all other regions >2 kbp from a gene locus. Of the 75,104 UMR-seq unique UMRs, 60,136 (80.1%) overlap an intergenic region compared to only 35,742 (38.9%) of the Shared UMRs. **d-e,** EM-seq normalised read-depth is visualised over Shared UMRs and UMR-seq unique UMRs. EM-seq read coverage is slightly reduced over UMR-seq unique, but highly similar to the coverage over Shared UMRs, and enriched relative to the methylated regions flanking the UMRs, suggesting that mappabillity of EM-seq reads is not significantly different at UMR-seq unique UMRs compared to Shared UMRs, as visualised for an example UMR-seq unique UMR in (**e**). **f,** EM-seq UMRs were re-called allowing regions containing only CHH sites (no CGs or CHGs), to be classified as EM-seq UMRs, then overlapped with UMR-seq UMRs. Compared to the earlier analysis, 20,010 UMR-seq unique UMRs were reclassified as Shared UMRs. **g,** The average q-score of UMRs identified by both EM-seq and UMR-seq ("Shared UMRs") is higher compared to UMR-seq UMRs that do not overlap EM-seq UMRs ("UMR-seq unique UMRs"). A lower q-score in epic2 indicates that the peak is less significant compared to the background signal, which would be expected of a partially methylated region that is only a UMR in a subset of cells leading to a relatively lower UMR-seq read signal. **h,** The average size and distribution of Shared UMRs are longer compared to UMR-seq unique UMRs. **i,** The majority of UMR-seq UMRs that are less than 300 bp (left panel) or less than 550 bp (right panel) do not overlap with EM-seq UMRs. Only 7.5% of the <300 bp UMR-seq UMRs overlap an EM-seq UMR; and only 19% of the <500 bp UMR-seq UMRs overlap an EM-seq UMR. **j,** Genome browser view of a UMR-seq unique UMR that is not detected by EM-seq because only ~200 bp is unmethylated, boarded by sparse methylation. **k,** Summary of the breakdown of UMR-seq uniques UMRs: of the 75,739 unique UMRs, 23,010 (20%) occur in regions containing only CHH sites, that can be reclassified as Shared by permitting these to be UMRs in the EM-seq pipeline; 19,563 UMR-seq UMRs are less than 300 bp in length, some of these also occur in CHH-only regions, with the remaining 14,766 accounting for another 20% of the UMR-seq unique UMRs. Based on the evidence in the heatmaps, metaplots and inspection of numerous individual loci, the remaining UMRs appear to be attributed to partial or sporadic methylation, meaning they failed to be called UMRs in the EM-seq pipeline.

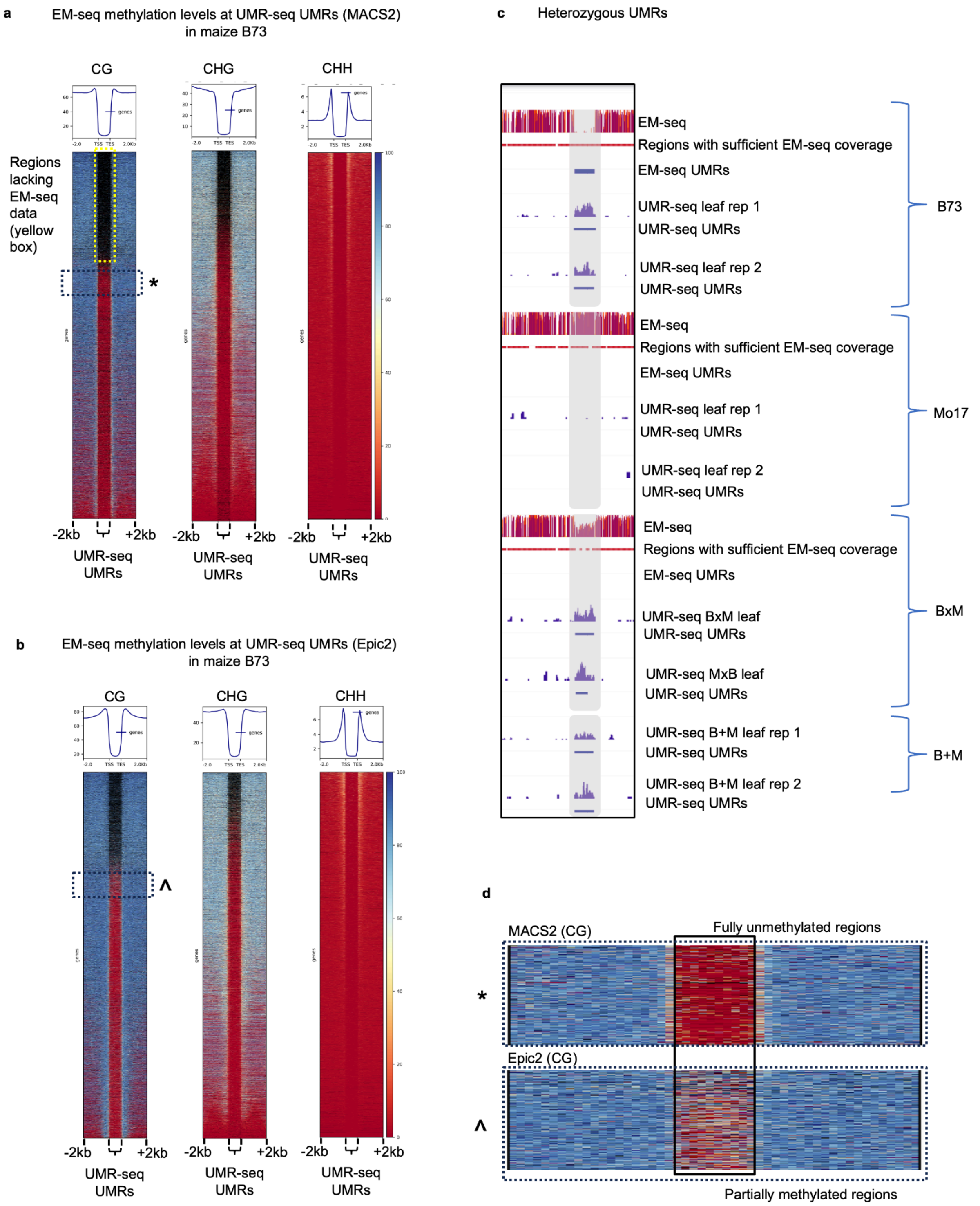


**Supplementary Figure 12. EM-seq methylation levels for UMRs identified using UMR-seq in maize.** Unmethylated regions (UMRs) were identified in UMR-seq data in maize using MACS2 in narrow peak mode **a,** or using epic2 in 50 bp sliding windows **b**, and excluding recalcitrant (blacklisted) regions. Using the same DNA sample, EM-seq libraries were prepared and sequenced and the DNA methylation levels for the UMR-seq UMRs were determined using the EM-seq data. DNA methylation levels are visualised for the UMR and 2kb either side. The black dashed boxes indicate the region magnified in panel (**d**). The yellow dashed box indicates UMRs with no data in EM-seq (due to the absence of a sufficient number of cytosines in that context or insufficient read coverage). **c**, Browser view of a UMR that is present in B73 but absent in Mo17. The UMR is then examined in a B73 x Mo17 F1 hybrid (BxM) and a Mo17 x B73 F1 hybrid (MxB), and also in a UMR-seq library that was prepared after mixing genomic DNA from B73 and Mo17 in a 50:50 ratio (B + M) to create an *in vitro* ‘hybrid’ sample to mimic a heterozygous state. In both the BxM and MxB hybrids and the *in vitro* mixed B+M samples, the heterozygous UMR can be detected using UMR-seq. Browser presents UMR-seq normalised read coverage and Epic2-called peaks regions. **d**, Magnified region of the heatmaps to compare methylation levels at MACS2 identified UMRs and epic2 identified UMRs. Epic2 has a greater tendency to identify partially methylated UMRs.

**a**

| *In silico* digestion modelling to identify recalcitrant regions (blacklisted regions) | | | | | | | | |
|---|---|---|---|---|---|---|---|---|
| Species | Plant family | Cotyledon No. | Ploidy level | Genome size (Mb) | No. of blacklisted regions | Total blacklisted regions (bp) | Blacklisted regions relative space to genome (%) | Estimated UMR-seq coverage (%) |
| Wheat | Poaceae | Monocot | 6 | 17,000 | 3,119 | 479,276 | 0.003 | 99.997 |
| Rice | Poaceae | Monocot | 2 | 430 | 228 | 30,456 | 0.007 | 99.993 |
| Maize | Poaceae | Monocot | 2 | 2,365 | 1,124 | 168,162 | 0.007 | 99.993 |
| Sorghum | Poaceae | Monocot | 2 | 772 | 1,010 | 172,501 | 0.022 | 99.978 |
| Barley | Poaceae | Monocot | 2 | 5,100 | 4,033 | 2,033,292 | 0.040 | 99.960 |
| Benth | Solanaceae | Dicot | 4 | 2,800 | 10,614 | 1,384,529 | 0.049 | 99.951 |
| Soybean | Fabaceae | Dicot | 2 | 978 | 4,256 | 514,642 | 0.053 | 99.947 |
| Grapevine | Vitaceae | Dicot | 2 | 500 | 4,059 | 489,829 | 0.098 | 99.902 |
| Tomato | Solanaceae | Dicot | 2 | 950 | 5,098 | 1,001,789 | 0.105 | 99.895 |

**b**

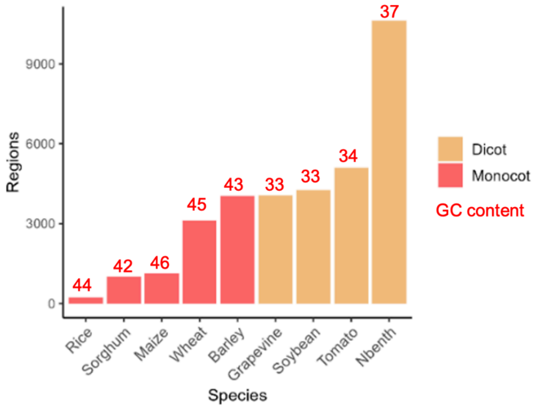


**Supplementary Figure 13. *In silico* digestion modeling across species a,** Genomes were digested *in silico* to identify DNA loci that would be recalcitrant to digestion by MspJI and/or FspEI due to the absence of the enzymes' recognition sequences. Numbers in the table represent sites that are recalcitrant to digestion into fragments smaller than 100bp, which could potentially lead to methylated DNA fragments larger than 100 bp being captured and sequenced during the UMR-seq library preparation process. **b,** The number of recalcitrant regions (blacklisted regions) were compared between monocots and dicots of varying genome sizes. Dicots have more recalcitrant regions.

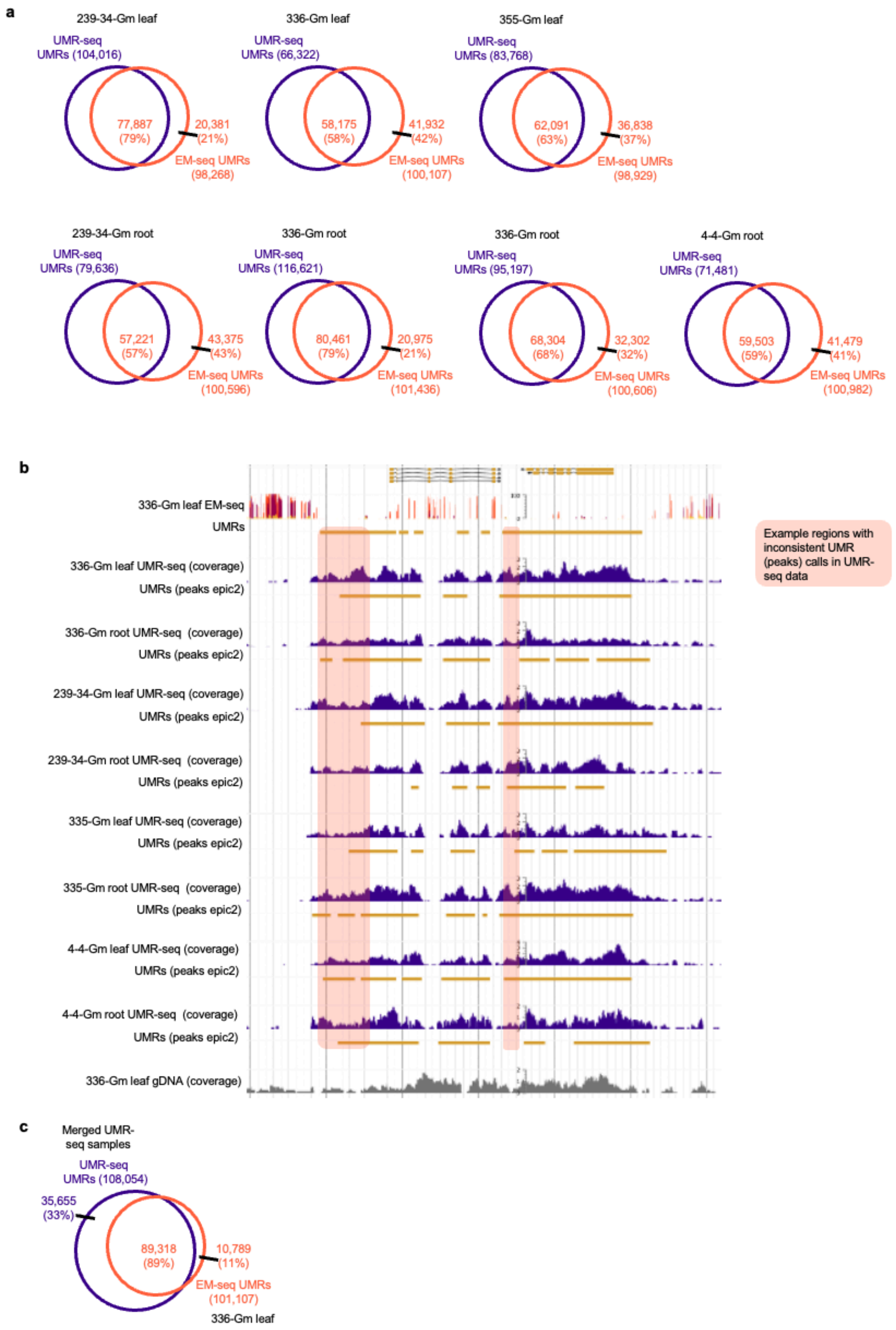
a
239-34-Gm leaf
UMR-seq UMRs (104,016)
77,887 (79%)
20,381 (21%)
EM-seq UMRs (98,268)
336-Gm leaf
UMR-seq UMRs (66,322)
58,175 (58%)
41,932 (42%)
EM-seq UMRs (100,107)
355-Gm leaf
UMR-seq UMRs (83,768)
62,091 (63%)
36,838 (37%)
EM-seq UMRs (98,929)
239-34-Gm root
UMR-seq UMRs (79,636)
57,221 (57%)
43,375 (43%)
EM-seq UMRs (100,596)
336-Gm root
UMR-seq UMRs (116,621)
80,461 (79%)
20,975 (21%)
EM-seq UMRs (101,436)
336-Gm root
UMR-seq UMRs (95,197)
68,304 (68%)
32,302 (32%)
EM-seq UMRs (100,606)
4-4-Gm root
UMR-seq UMRs (71,481)
59,503 (59%)
41,479 (41%)
EM-seq UMRs (100,982)
b
336-Gm leaf EM-seq
UMRs
336-Gm leaf UMR-seq (coverage)
UMRs (peaks epic2)
336-Gm root UMR-seq (coverage)
UMRs (peaks epic2)
239-34-Gm leaf UMR-seq (coverage)
UMRs (peaks epic2)
239-34-Gm root UMR-seq (coverage)
UMRs (peaks epic2)
335-Gm leaf UMR-seq (coverage)
UMRs (peaks epic2)
335-Gm root UMR-seq (coverage)
UMRs (peaks epic2)
4-4-Gm leaf UMR-seq (coverage)
UMRs (peaks epic2)
4-4-Gm root UMR-seq (coverage)
UMRs (peaks epic2)
336-Gm leaf gDNA (coverage)
Example regions with inconsistent UMR (peaks) calls in UMR-seq data
c
Merged UMR-seq samples
UMR-seq UMRs (108,054)
35,655 (33%)
89,318 (89%)
10,789 (11%)
EM-seq UMRs (101,107)
336-Gm leaf

**Supplementary Figure 14. Replicated UMR-seq in grapevine improves overlap with EM-seq UMRs. a**, Owing to the relatively high number of UMRs uniquely identified by EM-seq in grapevine compared to other species, we created 7 paired UMR-seq and EM-seq libraries from different genotypes and tissues. The reciprocal overlap between samples varied, with between 21-43% of EM-seq UMRs uniquely identified in EM-seq. **b**, When the UMR-seq coverage and peak annotations were compared across samples, and compared to a reference EM-seq library, an inconsistency in peak identification (pink boxes) was observed, despite the very similar coverage profiles of all UMR-seq samples. This inconsistency may be due to increased heterozygosity, reducing UMR signal which may affect EM-seq less if SNPs in the alternate haplotype/allele affect mapping to the alternative alleles. In this scenario, UMR-seq signal would be reduced relative to homozygous alleles, while EM-seq would not detect the alternative allele at all, thus calling a region an UMR regardless. Alternatively, the overall high level of unmethylated DNA in the grapevine genome leads to long stretches of UMRs, which may affect background estimation and UMR peak detection by algorithms such as epic2, increasing false negative rate in small lowly methylated genomes. This suggests that when undertaking comparative analysis of grapevine samples (or other small and lowly methylated genomes) using UMR-seq, non-peak based methods may be better suited to compare UMRs between samples. **c**, To investigate this further and to account for inconsistent per-sample peak identification, we merged the 8 available UMR-seq samples but merging together the peak files, which gave an improved overlap, leaving only 11% of the EM-seq UMRs, unique to EM-seq.

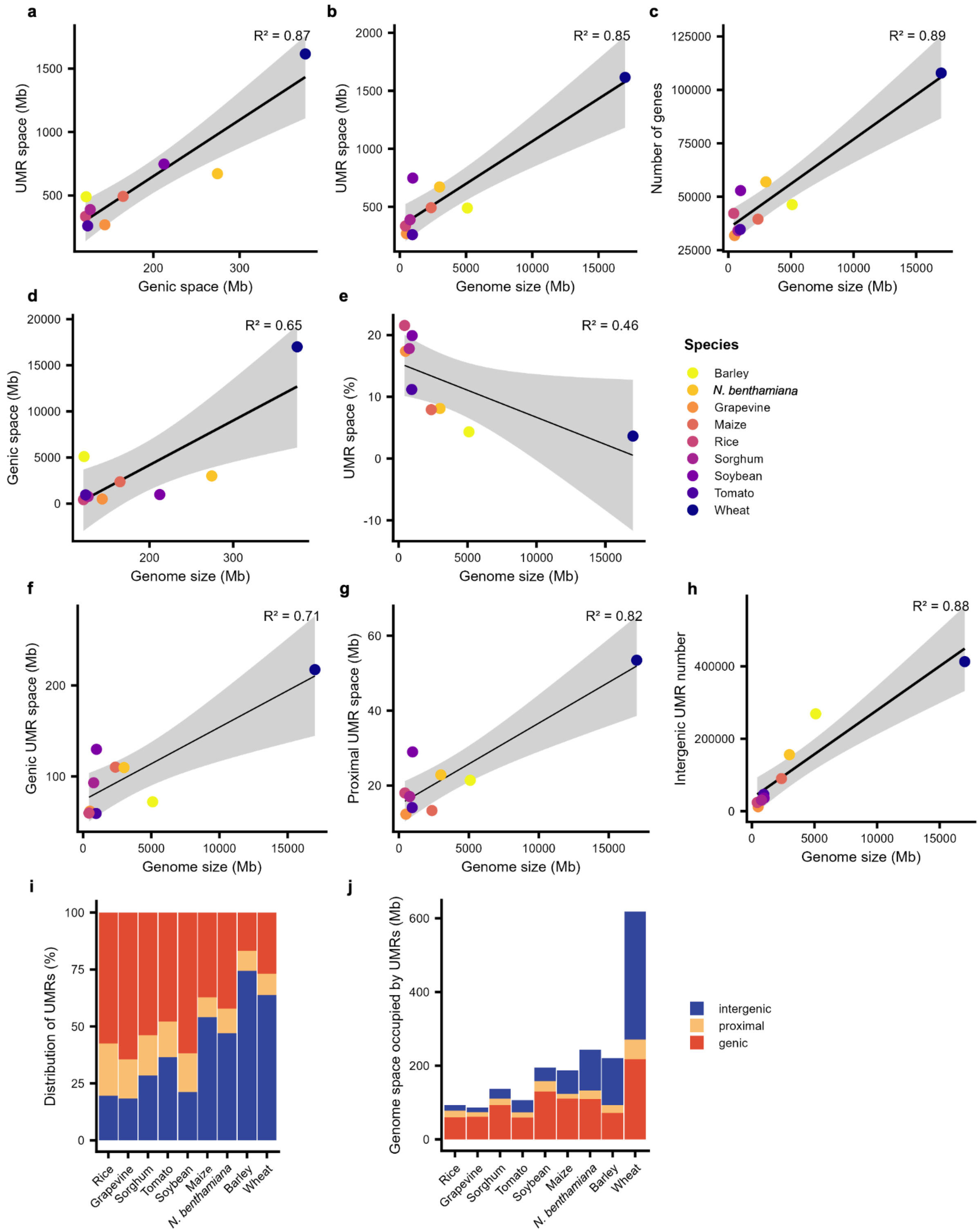


**Supplementary Figure 15. Relationships between unmethylated regions (UMR) space and genomic features across plant species. a - b**, Scatterplots show relationships between UMR space (Mbp) and genic space (Mbp) (**a**), and total genome size (Mbp) (**b**) across nine plant species. **c**, Shows the correlation between gene number and genome size (Mbp). **d**, Shows the relationship between genic space (Mbp) and genome size (Mbp). **e**, Shows the correlation between the proportion of UMR space relative to the total genome size (Mbp). **f** - **h**, Show the relationship of UMR space (Mbp) of each UMR subclass - genic (**f**), proximal (**g**) and intergenic (**h**) - with total genome size (Mbp). Each point represents a species, coloured according to the legend on the right. Solid black lines indicate linear regressions and shaded grey areas representing 95% confidence intervals.

$R^2$ is the coefficient of determination for the linear model. **i - j**, Bar plots showing (**i**) the distribution of all UMRs between subclasses (blue is intergenic, yellow is proximal, and red is genic), and (**j**) the portion of genome space (in Mbp) occupied by each UMR subclass across the nine plant species, organised by ascending genome size.

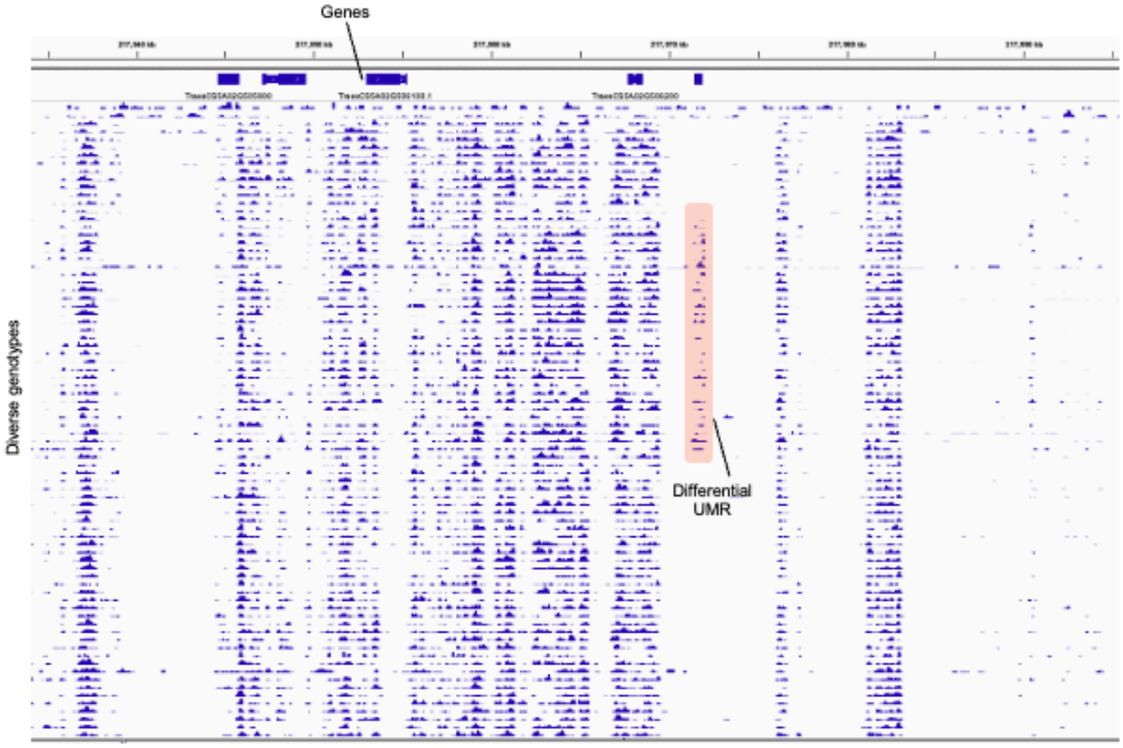


**Supplementary Figure 16. UMR-seq profiles of elite wheat accessions.** Example UMR-seq profiles of a sample of the wheat accessions, exhibiting an overall high similarity. Differential and presence/absence of UMRs also evident as highlighted by the pink box in the region of the Wheat Chinese Spring IWGSC RefSeq v1.0: chr5A_part2:134,173,007-134,211,502.

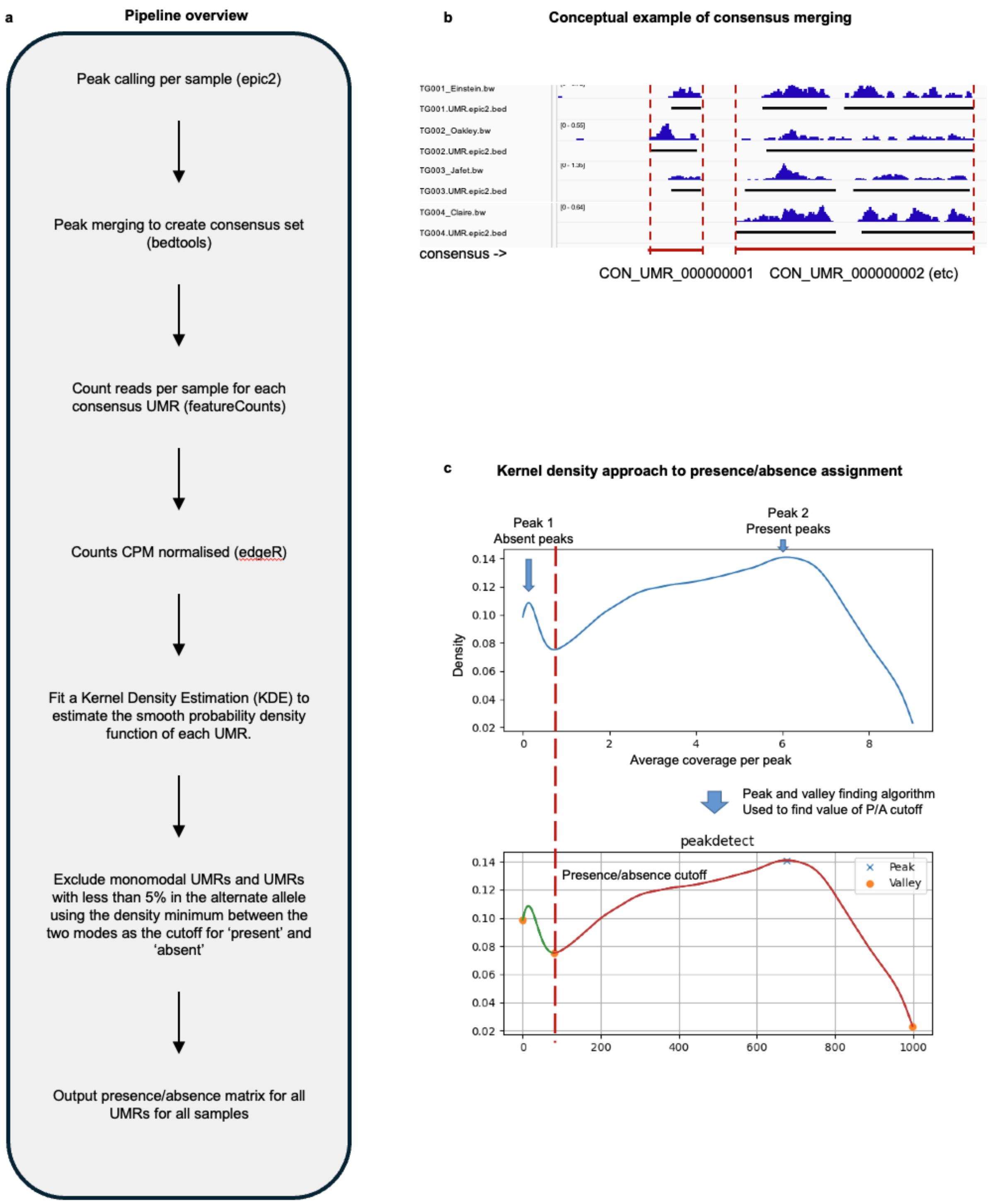


**Supplementary Figure 17. Consensus UMR pipeline for polymorphic UMR identification on the population scale a-b**, (**a**) Unmethylated regions (UMRs) were first identified in each sample with epic2 [46] (v0.0.41) in 50 bp bins, using Bowtie2 [84] alignments filtered to MAPQ ≥ 10 and deduplicated using samtools rmdup [96]. A unique identifier was assigned to every UMR in each sample. The UMRs from all samples were then pooled, sorted, and merged with bedtools merge [92] (v2.30.0), so that overlapping or book-ended UMRs from different samples formed a single locus (example in (**b**)). A consensus interval was then defined for each merged locus. UMRs more than ten times longer than the median length of the UMRs in that locus were excluded as outliers. The median start and median end coordinates of the remaining UMRs were taken as the consensus interval. Where this median interval was less than half the length of the full span of the locus (minimum start to maximum end), the full span was used as the consensus UMR and the median interval was recorded separately as an alternative interval. Alternative intervals were excluded from downstream analysis of the wheat panel in this analysis. Read support for each consensus UMR was quantified per sample with featureCounts [94] (v2.0.1). Only fragments meeting all of the following were counted: both reads of the pair aligned; the pair was not chimeric; the fragment length was between 50 and 1,000 bp; the mapping quality was $\geq 10$. The resulting UMR × sample count matrix was normalised to counts per million (CPM) with edgeR [95] (v4.4.2). Library sizes were defined as the total fragments assigned to consensus UMRs in each sample. **c,** Each consensus UMR was then called present or absent in each sample from the distribution of its CPM values across the cohort, using a kernel density approach developed by A. A. Golicz. *Density estimate*: The top 2% of CPM values for the UMR were removed to limit the influence of outliers. A univariate Gaussian kernel density estimate (KDE) was fitted to the remaining values with statsmodels [104] (v0.13.5), using a normal-reference bandwidth, and evaluated at 1,000 evenly spaced points across the observed range. *Peaks and minima*: Local maxima and minima of the density were identified with the peakdetect method in findpeaks [105] (v2.4.6), with a lookahead of 50 points. *Bimodality test*: A UMR was considered to have a bimodal distribution if the density had a single interior minimum separating one or two maxima. Where additional minima were detected, the UMR was also accepted if all maxima beyond the second interior minimum were lower than every maximum before it. UMRs that did not meet these criteria were not called. *Calls*: For bimodal UMRs, the first interior density minimum was used as the threshold. Samples with CPM above the threshold were called present and the remainder absent. *Final filter*: Calls were retained only if more than 5% of samples fell into each of the present and absent groups. This excluded UMRs where the apparent split was driven by a few outlying samples. To check the thresholds visually, KDE and peak/minimum plots were inspected for a random subset of called UMRs.

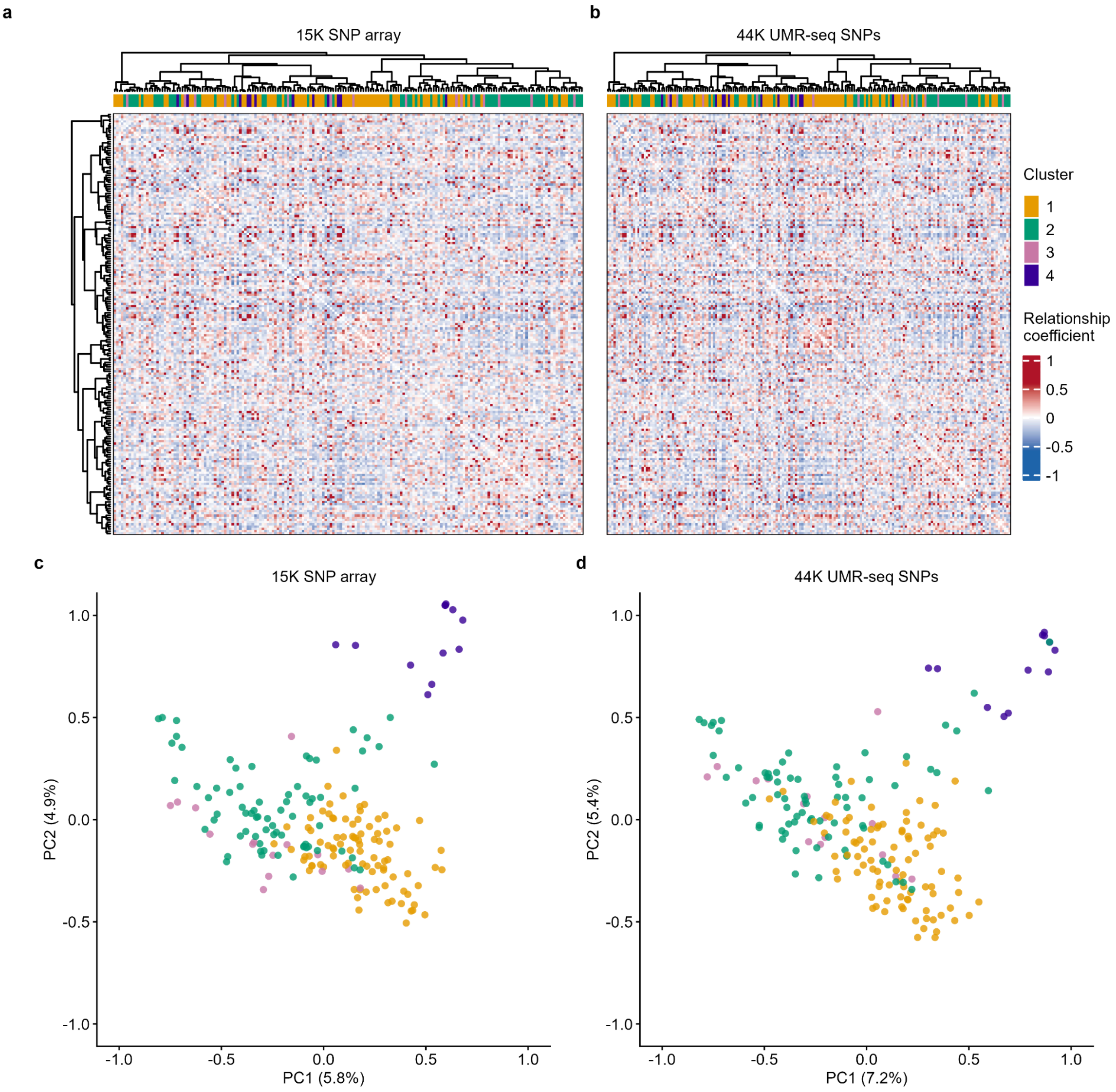


**Supplementary Figure. 18. Concordance between population structure inferred from UMR-seq SNPs and an independent SNP array.**

**a,b,** VanRaden relationship matrices for 187 European elite wheat lines estimated from the 15K SNP array (10,352 markers; a) and UMR-seq SNPs (43,622 markers; b). Matrices are ordered using a Ward.D2 dendrogram derived from the SNP-array matrix. The colour bars show the four groups defined from that matrix; the UMR-seq SNP matrix is therefore displayed in the order derived from the SNP-array matrix. **c,d,** The first two principal coordinates of the corresponding relationship matrices, coloured by the same groups and sign-aligned between panels. Off-diagonal matrix elements are correlated (Pearson $r = 0.87$).

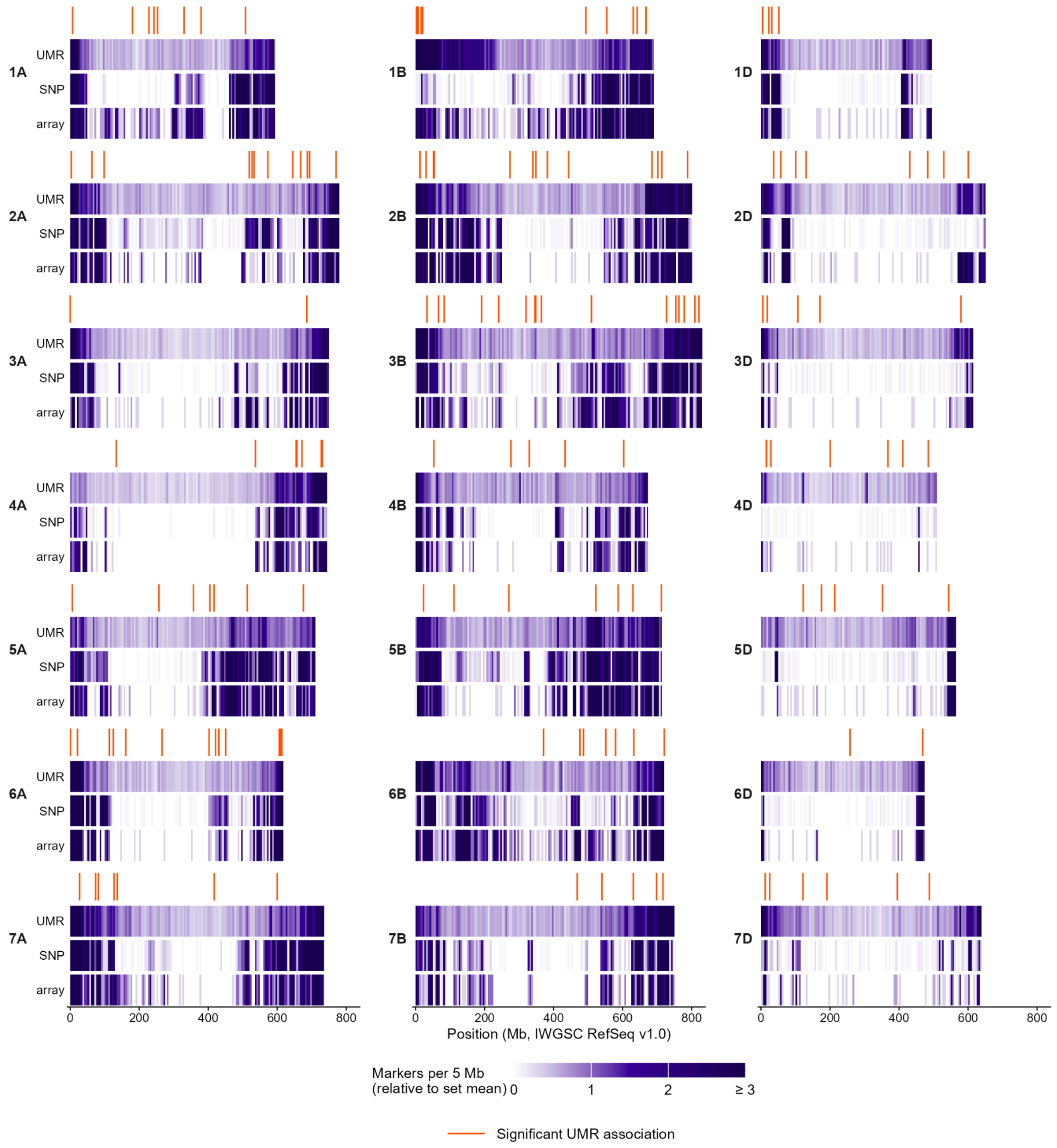


**Supplementary Figure. 19. Genome-wide distribution of UMR and SNP markers.**

Tracks show marker counts per 5 Mb window for UMRs (132,433), UMR-seq SNPs (42,905) and SNP-array markers (8,781 with chromosome positions) across the 21 wheat chromosomes (IWGSC RefSeq v1.0). Counts are expressed relative to the genome-wide mean for each marker set, with the colour scale truncated at 3. Ticks above each chromosome indicate the 183 distinct and significant UMR associations. Mean marker densities are 9.4 UMRs, 3.1 UMR-seq SNPs and 0.6 SNP-array markers per Mb. Every 5 Mb window contains at least one UMR, whereas 35% and 48% of windows contain no UMR-seq SNPs or SNP-array markers, respectively. In the D subgenome, UMR-seq SNP density is 1.1 per Mb, with 50% of windows empty.

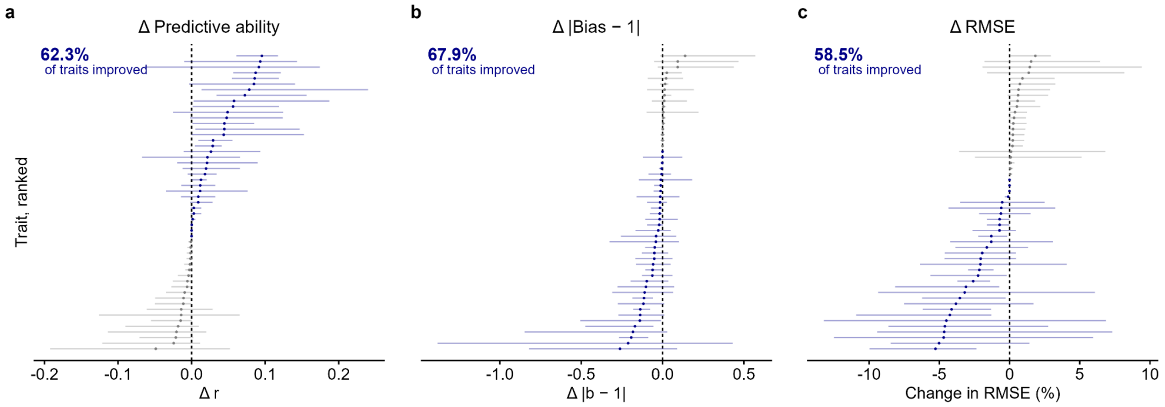


**Supplementary Figure 20. Changes in prediction performance after adding the UMR kernel.**

Prediction was evaluated across population groups for 53 trait–management combinations. Lines were assigned to five groups by partitioning around medoids using SNP relationships, and each group was predicted using the other four as the training set. **a**, Change in predictive ability (Δr). **b**, Change in the absolute deviation of the regression slope from one (Δ|b − 1|). **c**, Percentage change in root mean squared error. Points indicate the mean paired difference between models (SNP-plus-UMR minus SNP-only) for each trait–management combination. Combinations are ranked by their mean difference, and navy indicates improvement. Percentages report the proportion of combinations improved: 62.3%, 67.9% and 58.5% for **a–c**, respectively (Wilcoxon $P$ = 0.003, 0.0003 and 0.004).
